# Spontaneous rotations in epithelia as an interplay between cell polarity and boundaries

**DOI:** 10.1101/2021.11.11.468187

**Authors:** S. Lo Vecchio, O. Pertz, M. Szopos, L. Navoret, D Riveline

## Abstract

Directed flows of cells *in vivo* are essential in morphogenesis. They shape living matter in phenomena involving cell mechanics and regulations of the acto-myosin cytoskeleton. However the onset of coherent motion is still poorly understood. Here we show that coherence is associated with spontaneous alignments of cell polarity by designing cellular rings of controlled dimensions. A tug-of-war between polarities dictates the onset of coherence, as assessed by tracking live cellular shapes and motions in various experimental conditions. In addition, we identify an internally driven constraint set by cellular acto-myosin cables at boundaries as essential to ensure coherence, and active force is generated as evaluated by the high RhoA activity. The cables are required to trigger coherence as shown by our numerical simulations based on a novel Vicsek-type model including free active boundaries. We quantitatively reproduce *in silico* coherence onsets and we predict criteria leading to coherence. Altogether, spontaneous coherent motion results from basic competitions between cell orientations and active cables at boundaries.

## INTRODUCTION

Morphogenesis is driven by the interplay between Rho GTPases and the cytoskeleton ^1–4^. In particular, the acto-myosin network drives morphogenesis at the cellular level by elongation or neighbor exchange ^5,6^, and the cross-talk between Rho signaling and cellular mechanics has emerged as a generic property of morphogenetic systems. Interestingly, these dynamics are often associated with directed motions of cells. Directional flows in epithelia are reported during development in a variety of systems ranging from *Drosophila* to zebrafish ^7–9^. For example, during the development of spherical mammary acini or during the egg chamber elongation in *Drosophila*, cells exhibit *coherent* rotation on large scales with speed up to 20μm.h^-1 10–13^. These motions are proposed to be important for the robust morphogenesis of embryos.

The collective motion of cells is difficult to understand based on a simple analysis of migrations at the single cell level. Individually, cells can move randomly or directionally. However the epithelial layers which connect cells through adherens junctions may or may not coordinate the motion of the group ^14–16^. The understanding of emergence of collective effects requires theoretical approaches in tight comparison with experiments. In addition, although the interplay between RhoA activity and the acto-myosin cytoskeleton is documented at the single cell level ^1,17,18^, it is not yet fully characterized at the multi-cellular scale. In this context of potential collective effects, the understanding of coherent motion requires to study cellular assembly in simple geometries *in vitro* compared quantitatively to numerical simulations.

To this aim, we generated rings of epithelial cells, a geometry proposed by Turing in his 1952 article ^19^. This initial condition with boundaries is distinct from former work ^20,21^ and led to the formation of rings of connected cells, a simple and reproducible periodic configuration with no confinement. This allows the characterization of the spontaneous evolution of the cellular ring. We varied the ring diameter to identify the inherent coherence length of our epithelial MDCK cells. We report that below a threshold perimeter, rings undergo spontaneous rotations. We next sought to extract the ingredients generating this coherence. We show that an internal tug-of-war between cell polarities within the ring determines the onset of coherence, as shown by the initial polarity distributions and the time required to reach coherence. Tracking of cell division as polarity breaker supports this framework. We also report that coherence is associated with two supracellular acto-myosin cables at the inner and outer boundaries of the monolayer: these continuous structures act as autonomous confinements as they exhibit high RhoA activity and are shown to prevent cells from spreading out and breaking coherence. Finally, we test *in silico* our central mechanism of polarity tug-of-war coupled to the acto-myosin cables with a Vicsek-based model writing the fundamental laws of dynamics at cellular levels for polarity, velocity and including forces at boundaries. After calibration, we show that this computational approach predicts the dynamics of all rings with no ad hoc adjustments. We also calculate phase diagrams for coherence as a function of cellular parameters combinations. Our results show that activity at boundaries – supported by RhoA activity in experiments – is indispensable to ensure cellular coherence. We propose that cell polarity tug-of-war together with self-generated confinement from spontaneously assembled active cables set the tempo for coherent movements in epithelial monolayers.

## RESULTS

### Obtaining coherent motion

We first needed to identify the natural length over which cells migrate directionally – namely the maximal coherence length *ξ*_*max*_– characterizing our system. A circular array of cells is expected to undergo rotation if the maximal coherence length *ξ*_*max*_ matches the perimeter p. If the coherence length is smaller than the perimeter, non-coherent motion is expected ^21,22^. To obtain *ξ*_*max*_, we prepared multicellular rings with free boundaries and decreasing diameters by using micro-contact printing fibronectin regions and differential adhesion (see Methods) : 1000μm, 300μm and 180μm (see Movies S1 to S3, respectively), and we measured the 12-hours velocity fields (see Methods). Without confinement, cells could spread out on passivated and non-coated regions, and rings had different behaviors: large rings – 1000μm – exhibited growth of cellular fingers inwards and outwards whereas smaller rings – 300μm and 180μm – behaved like cellular assemblies during wound healing with centripetal closure (see Fig.1a and Movies S1-S3). For 1000μm and 300μm rings, we saw local coherent flows of length *ξ* (Fig. 1b) but no global coherence. However, at 180μm diameter, about a quarter of rings spontaneously rotated with a velocity of ∼ 20μm.h^-1^, which suggested that we approached the perimeter value potentially leading to coherent motion. To find this perimeter, we plotted the correlation function of tangential velocity, and we extracted the maximal coherence length *ξ*_*max*_(see Fig.1c, Fig.S1a and Methods) ^23^. We show that *ξ*_*max*_, obtained from the x-intersect of the tangent at origin of the correlation function, is similar in every condition – *ξ*_*max*_ = 315μm (Fig. 1c and Fig. S1a) –, demonstrating that *ξ*_*max*_ is an inherent length of our cellular system.

**Figure 1.**
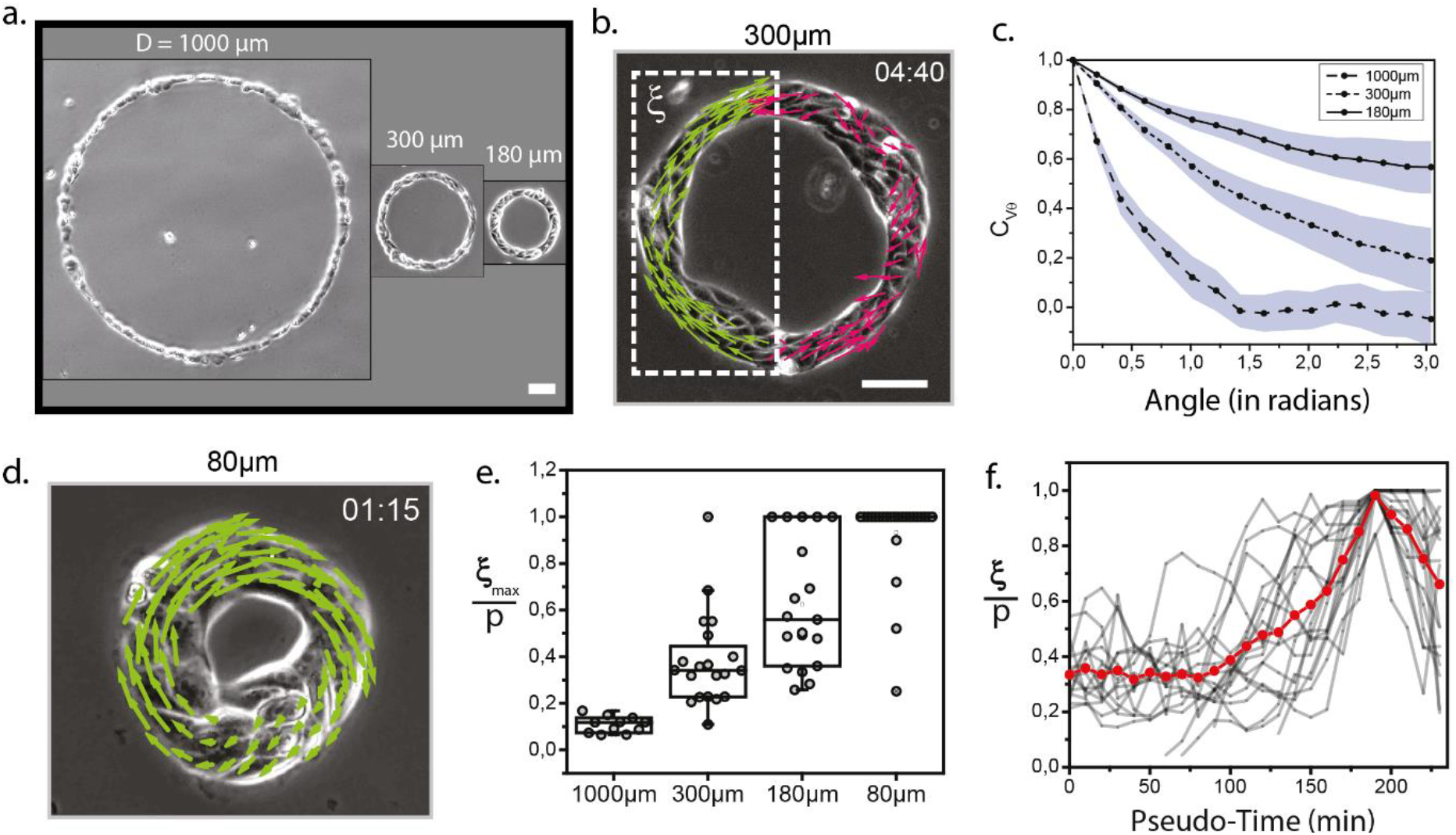
Acquisition of coherent motion. (a) MDCK rings of various diameters at initial time t_0_. (b) Coherent flows can be extracted from the velocity fields. A coherent flow of length ξ is highlighted in green. Time in hh:mm; Scale bar = 50μm. (c) Average correlation functions of tangential velocity v_θ_ for each ring diameter. The angle θ corresponds to the polar coordinate on the ring. (d) 80μm ring with a perimeter below the coherence length undergoes spontaneous rotation. (e) Index of global coherence 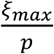 for each ring diameter. 80μm rings are “rectified” and the majority of rings are coherent 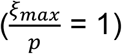. n_1000_ = 11 ; n_300_ = 20 ; n_180_ = 20 ; n_80_ = 24. Medians are shown as lines in the boxplot. (f) Temporal sequences are aligned on the coherence acquisition 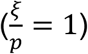 along a new pseudo-time. The red line corresponds to the mean of all experiments. Coherence is a non-linear process taking place within ∼1h30.

Based on this *ξ*_*max*_ of 315μm, we designed and focused on rings with 80μm diameter corresponding to 250μm perimeter below *ξ*_*max*_. As expected, we “rectified” the motion of cells around the ring and we ended up with 83% of rings spontaneously rotating at a velocity of 20-25 μm.h^-1^ (see Movie S4, Fig. 1d and Fig. S1b). All results are then reported in Figure 1e, where 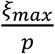, the coherence length over the perimeter, is defined as an index of global coherence: it is equal to 1 when the whole ring rotates and gets close to 0 as the global coherence decreases. Interestingly, there is a bias in clockwise rotation which suggests a spontaneous cellular bias as reported in previous works ^24,25^ (Fig. S1c).

If coherence was obtained for this 80μm ring diameter, the time needed to reach this complete rotation (defined as 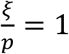, see Materials and Methods) varied from 1 hour to 10 hours and lasted from 1 hour to 10 hours (see Fig. S1d). This large distribution impeded quantification of the transition from non-coherent to coherent motions. Therefore, we needed to rescale the time needed to reach coherence starting from the initial condition of continuous ring of cells. We decided to align all plots with respect to the onset of coherence defined as 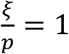. We could then determine the quantitative behavior of coherence acquisition. This pseudo-time representation allowed to show that coherence acquisition is a process lasting about 1h30 with minor variability among rings (Fig. 1f). This timescale is fast considering the 12-hours duration of the experiment and this can be viewed as a quick switch between the two states. At coherence, a continuum of behaviours appeared among the rings populations from rings with persistent rotation over hours to rings with transient coherent motion shorter than one hour (Fig. 1f and Fig. S1d-e). To understand the origin of these dynamics, we measured next the ring velocity and persistence of rotation, and saw their correlations (Fig. S1f). This suggests that collective cell speed might be involved in maintaining coherence even though inertia is negligible at this scale.

### Tug-of-war between cell polarities sets coherence

Next we sought to identify the cellular mechanism at play for the onset of coherent rotation. Cell polarity is a natural readout for its key role in setting direction of motion. We thus looked at the distribution of cell polarities inside rings. For this, we used labeled lamellipodia (see Fig. 2a, see Methods for the measurement of polarity). We observed selected lamellipodia orientation and direction. Initial polarities showed a bi-modal distribution peaked around tangential directions with two main orientations, *i*.*e*. 0° and 180° (see Fig. 2b). These two states suggest that a tug-of-war between polarities could happen between cells of opposite directions within the same ring. We reasoned that initial polarities could then play a role in the time needed to reach coherent rotation. A large number of cells with the same polarities would minimize the tug-of-war, and would decrease the time to coherence. We then followed each cell polarity defined by tight-junction organisation (see Fig. 2c and Movie S5). We had checked that the direction of shape anisotropy defined by cell geometry correlates with the direction of the lamellipodia (Fig. S2Aa-b). Our measurements of percentage of cells with aligned polarities as a function of time to coherence show a decreasing relationship (Fig. 2d). This further indicates that distribution of single cell polarities imposes a tug-of-war and determines the time needed to enter coherence. We tested this statement by inhibiting lamellipodia formation with the Arp2/3 inhibitor CK666 and this prevented coherence (Fig. S2B) by suppressing the tug-of-war between polarities.

**Figure 2.**
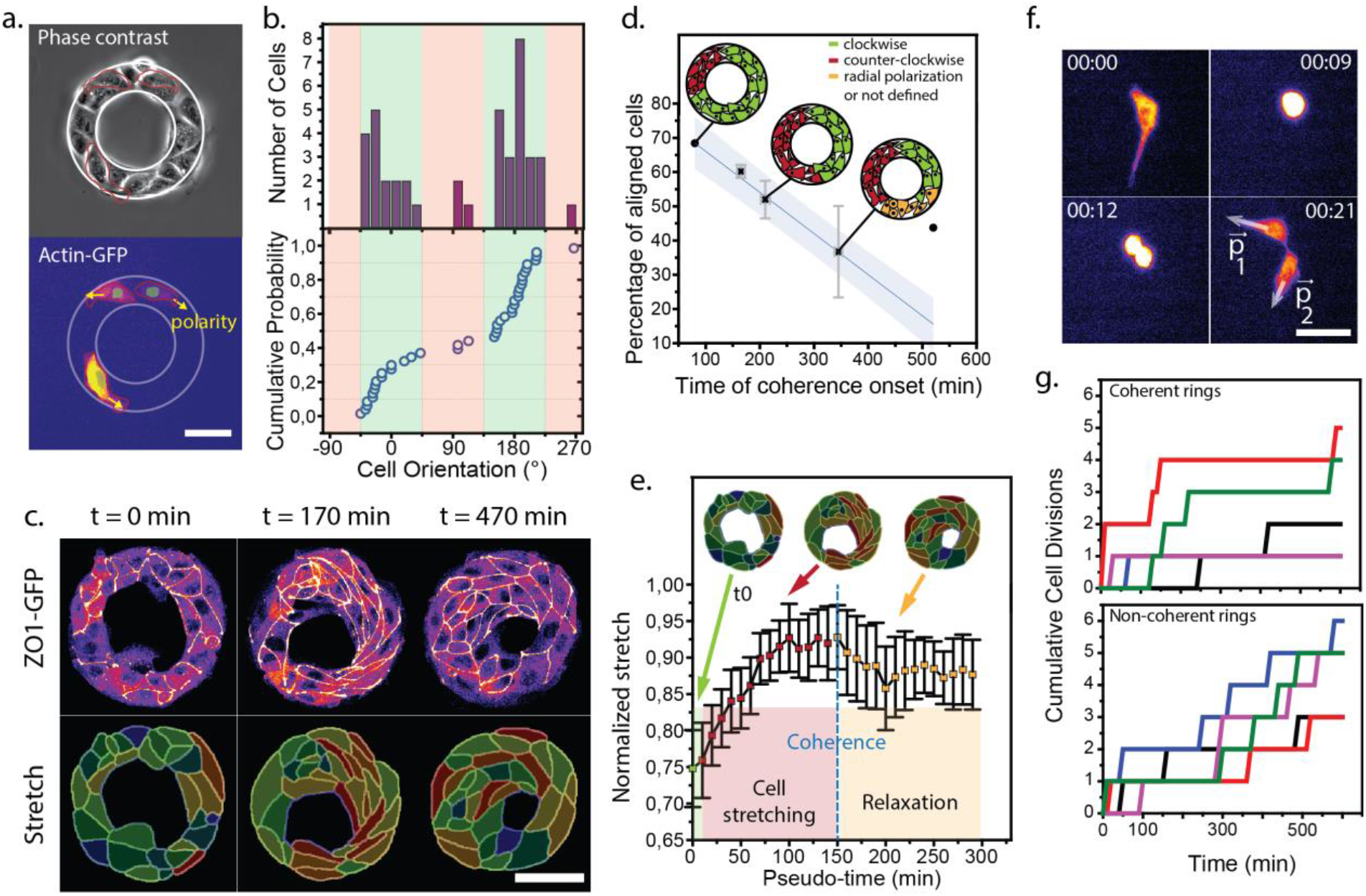
Cell polarity and density determine dynamics of coherent motion. (a) 80μm “mosaic” ring. Arrows indicate cell polarity based on the direction/orientation of cell lamellipodium. Scale bar = 50μm. (b) Distribution of cell initial orientations within the rings. 0° (counter clockwise) and 180° (clockwise) correspond to tangential orientations. n_rings_ = 12, n_cells_ = 42. (c) Segmentation of multicellular rings based on ZO1 distributions. Single cell parameters such as cell elongation (stretch) can be extracted, red and blue largest and lowest stretch respectively. Scale bar = 50μm. (d) Percentage of aligned cells based on the polarity given by ZO1 distributions. Data are binned with respect to the time of coherence onset. Data are represented as Mean ± SD. N = 10. (e) Average normalized stretch (initial/max) aligned along a new pseudo-time. The dashed blue line corresponds to coherence acquisition. Data are represented as Mean ± SD. N = 10. (f) Division of an actin-GFP labeled cell within a 80μm ring. Cell division generates daughter cells with opposite polarities **p**_**1**_ and **p**_**2**_. Scale bar = 20μm. (g) Cumulative cell divisions as a function of time. Coherent rings exhibit bursts of divisions with large periods without divisions whereas non-coherent rings undergo regular divisions throughout the course of the experiment. For clarity, only 5 representative rings are shown for each case. Full population is available in Fig. S2A.

The co-existence of opposite polarities is expected to generate stretching within the ring. From tight junctions contours, we could also extract cell elongation (Fig. 2c and Fig. S2A). Using the pseudo-time as the reference (Fig. 2e), we report three phases: (i) 2.5 hours before coherence, cells elongate until (ii) their tangential stretch reach constant values sustained for about 1 hour; (iii) coherence starts shortly within 1h followed by a rapid drop in elongation (see Fig. S2). This is distinct from non coherent rings (see Fig. S2Ac). These measurements substantiate our framework: cell lamellipodia “pull” in opposite directions leading to cellular stretch and resolution of this competition induces cell stretch relaxation visualized by the rapid drop in stretch.

### Cell divisions as polarity breakers impede coherence

To further test the central role for polarity in setting coherence, we tracked cell division events within rings. Right after mitosis, cells have opposite polarities and thus are expected to challenge polarity alignments. These opposite polarities indeed happened at cell division within rings (see Figure 2f). We quantified the effect by plotting the cumulative number of cell divisions within rings. We saw that rings where division regularly occurred were related to poorly coherent motion (Fig. 2g, bottom). In contrast, when a burst of many cell divisions occurred within a short time period (typically 2 hours over a total time of experiment of 12 hours), coherence was maintained in its optimal value (Fig. 2g, top). Divisions generate cells with opposite polarities and impede coherence (see Fig. S2Ae,g,h). Division as a polarity breaker acts as a coherence breaker.

### Acto-myosin cables as internally self-assembled boundaries imposing tangential cell polarity

So far, we reported that cells were polarized within the ring and we next sought to further understand why tangential orientation was selected (Fig. 2a-b). If the rings were confined by physical walls, this orientation would be favored^20^. However, in our assay, cells were free to move in any directions suggesting that an additional phenomenon biases cell polarities. We first stained for actin and myosin since acto-myosin cables were reported to self-assemble within cells at epithelial colony boundaries ^17,26,27^ (Fig. S3Aa). This generic feature was therefore expected to appear as well in our rings and we hypothesized that these structures could contribute to orient cell polarity. We show in Figure 3a the presence of these acto-myosin cables and we propose that cells internally set their “walls” which in turn could potentially make them be oriented along the perimeter rather than radially.

**Figure 3.**
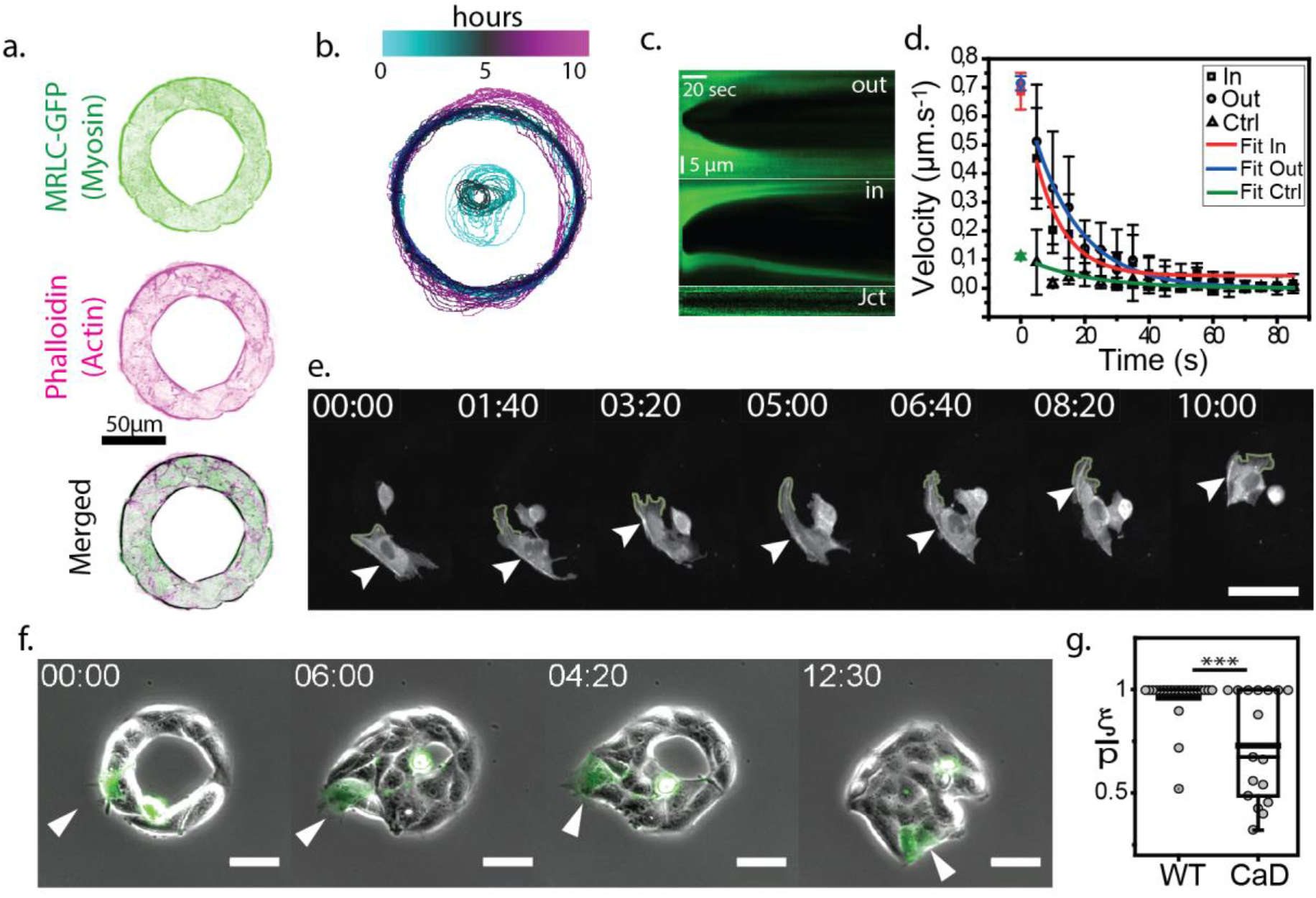
Acto-myosin cables as internally-driven constraints. (a) Immunostaining of a 80μm ring. Myosin and actin are visualized. (b) Tracking of the ring boundaries. The color code indicates time. The ring first closes inwards before extending outwards. (c) Kymographs showing acto-myosin ring retraction after laser ablation and cell-cell junction opening (as a control) after ablation. (d) Retraction velocity as a function of time after laser ablation. Exponential fits give estimates of v_0_ defining the relative tension stored in the ring. (e) Actin vizualisation (lifeAct) of a transfected cell within a 80μm ring. The acto-myosin cable is self-generated by the cell during motion and constrains the polarity in a tangential orientation. White arrow indicates the cable, green contour underlines the lamellipodium boundary. Scale bar = 20μm. (f) Time-lapse showing a CaD transfected cell (white arrow) escaping the multicellular ring and challenging coherence. Scale bar = 50μm. (g) Coherence acquisition for non-transfected ring and rings with CaD transfected cells. There is a significant decrease in coherence (p = 7.10^−4^) in the CaD rings.

To test this idea, we evaluated the mechanical contributions of these cables and their roles in cell confinement. First with live observations, we saw that the inner cable was contractile and led first to the closure of rings within hours prior to extension ‘out’ of the cellular disk now formed (Fig. 3b, Fig. S3Ab and Movies S3-4). This suggests that indeed inner cable pulls inwards as expected for a contracting acto-myosin cable. This dynamic was distinct from larger rings of 1mm (see Fig. 1 and Movie S1) which suggests that cable curvature plays a role in cell motion ^28^. This curvature dependence was further substantiated by experiments with elliptical geometries (see Fig. S3B). Then, we reasoned that the outer ring may act the same way by pushing cells inwards. To evaluate whether these structures were contractile, we used laser ablation experiments ^29^. First we observed a fast and large opening of both cables within seconds with a similar extent of ∼ 12μm (see Figure 3c and 3d and Movie S6, see also Fig. SA3b, and Methods). This demonstrates that cables are more tensile than cell-cell junctions also tested in these experiments which yielded small openings (∼ 1μm). In addition, we measured the opening velocity after laser ablation as a read-out for comparing stress between inner and outer cables assuming they bear the same damping coefficients ^29^. Our results suggest that acto-myosin cables indeed apply contractile constraints on the boundary cells and with similar values (see Fig. 3d).

Next, we tested the impact of this contraction at boundaries on cell polarity. We followed cells transfected with Lifeact within non labeled rings. We report in Fig. 3e and Movie S7 the interplay between cell polarity and the outer acto-myosin cable: cells could build their own acto-myosin cables while the structure imposes lamellipodia to be oriented tangentially. This supports the notion that acto-myosin cables and tangential polarity are correlated.

To further test this coupling, we sought to remove locally the influence of acto-myosin cables. We included within rings single cells expressing caldesmon (CaD) (Fig. 3f), a calmodulin binding protein inhibiting Myosin ATPase ^30^. Indeed this triggered a local decrease in myosin activity and in cable contractility. These caldesmon expressing cells extended out of the rings with a radial motion (Movie S8). This further confirms the key role played by the cables in confining cell polarity tangentially. This perturbation also impaired rotations in the majority of cases (see Fig. 3g and Suppl. Fig. 3d). On average, we found that coherence was lower on CaD mosaic rings (Fig. 3g) and this was correlated to the amount of CaD levels (Fig. S3Ae). Altogether, this suggests that continuity of the acto-myosin rings is important for tangential motion and coherence. In conclusion, cables are essential to confine and orient cells polarities tangentially and further trigger rotations.

### Force is active at boundaries and cell velocity is corelated with RhoA level

Observations and measurements on cell dynamics could hide signaling effects associated to regulations of the acto-myosin cytoskeleton by the small GTPase RhoA^31,32^. To evaluate this potential contribution, we analysed the correlation between spatio-temporal dynamics of RhoA activity and cell dynamics. For this, we used a FRET sensor which shows distributions of active RhoA in space and time (Fig. 4a and Ref.^18,33^, see Methods and Fig. S4a-c). In MDCK rings, we found the largest activity at inner and outer boundaries, where acto-myosin cables are assembled (Movie S9, Fig. 4b and Fig. 3a). This supports the contractile nature of these fibers. Accordingly, we found that the closure time of rings is shorter for large FRET activities in cables (see Fig. S4d-e). We can conclude that regulation by RhoA mediates active forces at boundaries.

**Figure 4.**
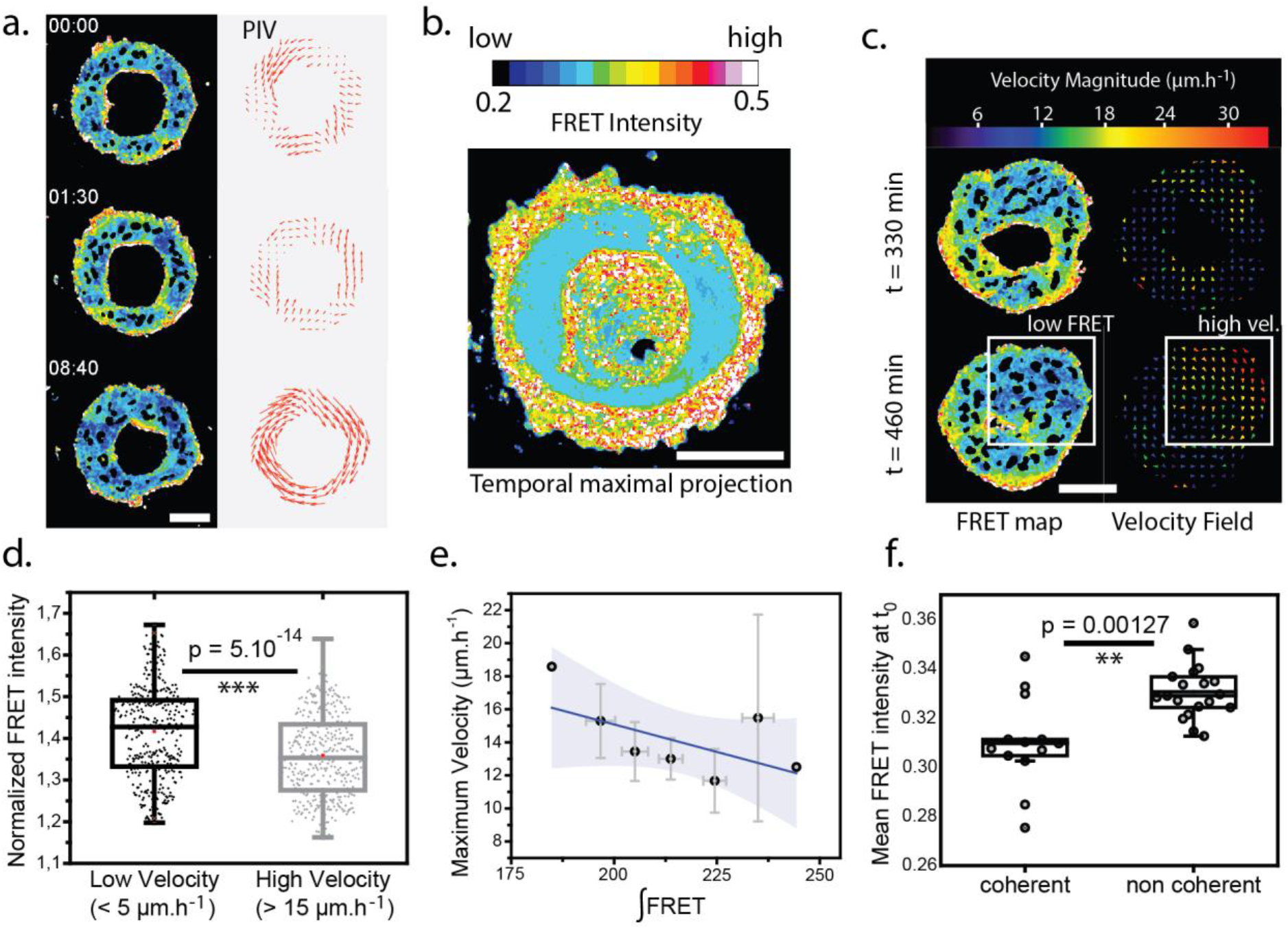
RhoA activity correlates with different ring behaviors. (a) RhoA activity on a 80μm ring as described by the FRET ratiometric map (left) associated to a velocity field (right). Scale bar = 50μm. (b) Temporal maximal projection showing highest FRET signals at ring boundaries. Scale bar = 50μm. (c) FRET map for each frame is represented with its associated velocity field. Scale bar = 50μm. Comparison is shown in (d) where high FRET regions correlate with low velocity. (e) Maximum velocity of the ring (with both radial and tangential components) is represented as a function of the averaged FRET level integrated over the entire experiment. (f) FRET intensity at t_0_ averaged over the cellular “bulk” excluding cables. Initial levels of RhoA activity are plotted for different coherent behaviors. High activity is correlated with low coherence. Data in (e) are binned; full distributions are shown in Fig. S4h. Same color codes for (a), (b) and (c) for FRET signals.

Building on this result, we sought to test whether correlation between RhoA and cell velocity could be at play in the onset of coherence, knowing that cell velocity plays a key role in collective cell motion. We analyzed locally and globally velocity fields and their associated FRET maps (Fig. 4c and Fig. S4h, f, g, see also Methods). We found that high velocity regions for cells were correlated with low FRET levels (Fig. 4d,e and Suppl Mat). This suggests an inverse relationship between the two quantities (see also Fig S4i). Cells move faster with lower levels of FRET. This further supports the notion that RhoA activity is essentially encoded in cell velocity.

We then turned to numerical simulations to check whether the onset of collective motions could happen with basic interactions rules extrapolated from our experimental results, i.e. active force at boundaries and cell velocity rule.

### Numerical simulations reproduce the ring dynamics qualitatively and quantitatively

We designed numerical simulations based on an original Vicsek type model with active boundaries, involving parameters accessible to experiments (see Fig. 5, Fig. S5, Table 1 and Annex_Math). Cells are represented as particles with velocity **v** and polarity **p**. Backed up by experimental observations (Fig. S5a-b), we assume that cell velocity aligns to its polarity. Cell polarity is allowed to diffuse with a coefficient D (Fig. S5d-e) and aligns with the average polarity around the ring with rate μ (Fig. S5f). Cell polarity is also assumed to align with cell velocity direction with rate ν. Local springs between boundary cells dynamically describe the inner and outer cables. Active and passive contributions of these cables are incorporated in the polarity dynamics. Table 1 summarizes data taken from experiments and inserted as an input to the model. More details are provided in Annex Math.

**Figure 5.**
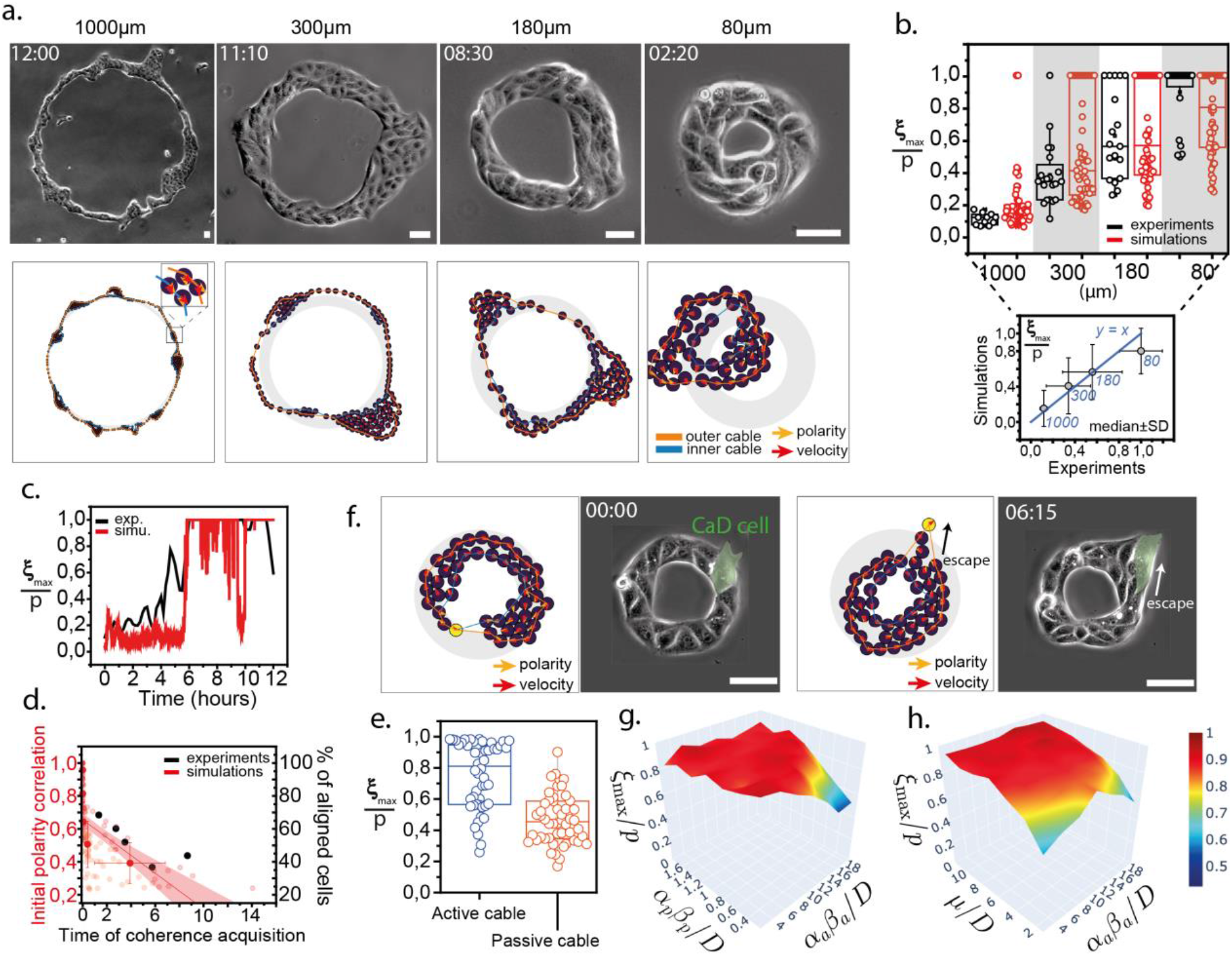
Theoretical modelling of multicellular rings. Experiments are compared with simulations. (a) Rings of different diameters are represented (top) when typical motifs appear, fingering for large diameters, clusters for intermediate rings, closure for smaller rings. Morphology is reproduced by numerical simulations (bottom). Scale bar = 50μm. Time in hh:mm; simulations times (in hh:mm, from left to right): 05:24; 17:56; 04:00; 04:57. (b) Coherence index as a function of ring diameters for simulations and experiments, with an inset comparing median values. (c) Dynamics of coherence acquisition *in silico* compared with a typical MDCK ring. (d) Initial polarity correlation as a function of the time of coherence acquisition. The decay observed in experiments is similar to simulated rings. (e) Coherence index for two different populations of MDCK rings: *activity* within cable facilitates acquisition of coherence. (f) Caldesmon experiment and cellular ‘escape’ are reproduced with simulations. The yellow particle represents the CaD cell and its cable parameters are set to zero. Scale bar = 50μm. Time in hh:mm. Simulation times (in hh:mm, from top to bottom): 05:20; 13:03. (g-h) *Phase diagrams for coherence with respect to the dimensionless parameters associated with the model (active cable force - active alignment, active cable force – passive cable force). Index* p *refers to the passive component of the cable while* a *corresponds to the active one. On each plot, each point of the surface corresponds to the average value of 50 simulations*.

We report in Fig. 5 the comparisons in shapes and coherence values across conditions. The ring phenotypes are reproduced with multicellular fingers or clusters for distinct colony diameters (Fig. 5a and Movie S10-13). Similarly, the coherence index is quantitatively obtained with excellent precision (see Fig. 5b) together with its typical time dependence (Fig. 5c). Also, the tug-of-war between polarities correlated with the time required to reach coherence for simulated rings with quantitative consistency with experimental times (Fig. 5d). We also found that closure time depends on the initial cell alignment (Fig. S6a).

In addition, we could probe numerically the potential contribution of active forces at cables: in our model, cables have a passive component corresponding to the stretching and bending moduli of the acto-myosin structure, and an active component translating the contribution of cables in setting tangential polarities (as explained above and shown in Figure 3). Interestingly, the active component was required for the onset of coherence (Fig. 5e). We also tested the effect of having a cell with no cable within the ring, thereby modelling the caldesmon experiment (Fig. 3f): the phenotype of this cell escape was reproduced as well (Fig. 5f and Movie S14), substantiating the confinement role of cables. We checked that extremely stiff cable acted as walls like in Jain et al. and led to coherent motion (Fig. A10 and A11, Annex Math).

Finally, we generated phase diagrams to understand the relationship between coherence and cell polarity and cables activity. We found the range of values necessary to trigger coherence for the main parameters of the model (cell polarity alignment, cable activity and physical properties) (Fig. 5g and 5h). We show that the coherence value is essentially governed by the cell polarity alignment: a large collective alignment increases coherence (Fig. 5g and Fig. S6c-d) while the bending and the stretching moduli (passive terms) do not affect coherence initiation at least in the range of considered parameters (Fig. 5h and Fig. S6b). However, the cables properties have a strong impact on the closure time of the ring and passive and active terms have antagonist effects in this regard (Fig. S5g and S5h).

Altogether our numerical experiments based on measured cellular parameters and with basic interactions for single cells show that we can capture the main features for the onset of coherence with no need to incorporate parameters other than polarity and active cables at boundaries.

## DISCUSSION

We show that coherence of cellular movements emerges when the system size is below the coherence length inherent to MDCK cells as reported previously on epithelial disks ^21,22,34,35^. Cells align their polarities within the ring and this step sets the onset of coherence. Furthermore, acto-myosin cables act as internally driven constrains confining cells tangentially. RhoA activity is larger at boundaries co-localizing with cables and is inversely related to the cell maximum velocity. Finally, with these ingredients, experimental and numerical rings share qualitative and quantitative features, supporting the relevance of our minimal framework for cell interactions. We show that active forces at cables are critical to orient cell polarity and contribute to coherence.

### Internally driven coherence versus externally driven coherence

Emergence of coherent flows has been studied *in vitro* in confined situations ^20–22^. Cells were chemically or mechanically confined and cannot escape their adhesive patterns ^21,22^. This led to distinct phenomena – although with similar velocities of 20 μm.h^-1^ - where the coherence is not driven only by the cellular characteristics but also by the boundaries of confinement as suggested by previous theoretical works ^36^. In Ref. ^20^, this led to rotations even on rings with a diameter of 1mm, far above the natural coherence length of the epithelial cell line. We reproduced this result numerically (see Sec. 3.4 Annex Math) which further shows that the external walls may impose the rotation for large scales. Intuitively this can be understood by the fact that cell motion within a cohesive ring monolayer can only be tangential with solid boundary. In our case, we let cells spontaneously self-organize without confinement and we decreased the pattern size to initiate the onset of coherence. This new configuration makes this rotation spontaneous in essence and not guided by 3D walls which force the motion along the ring. Remarkably, the symmetry of rotation clockwise-counterclockwise is broken on average, in contrast to former studies in confinement^20^ but in agreement with the bias reported in other studies^24,25,37^ with potential roles for left-right asymmetry during development. We make the hypothesis that chirality at the single cell level – generated by molecular structures with broken symmetry^25^ – is conserved through scales and is translated in the emerging collective motion ^38^. Altogether, our spontaneous broken symmetry reveals inherent cellular properties which convey spatio-temporal order selected by cells.

### Coherence length and spatial scales

It is interesting to analyze the potential origin of this natural coherence length. This length is distinct from former works reporting spontaneous flows of active nematics: rotation was not associated with opposite senses of rotation at boundaries, and the coherence length is higher compared to these 2D confluent monolayers. Here, cell motions are not impeded by cellular jamming seen in fully confluent monolayers^39^. Therefore, we propose that the coherence length found in our study is not expected to be the same as that defining the crossover from rotation to active turbulence seen in fully two dimensional confluent layers of cells.

In addition, the passage from single cell to tens of cells forming a new entity necessarily go through adhesion mediated by cell-cell junctions and its interplay with cytoskeleton. We indeed saw that rings with MDCK cells overexpressing cadherins presented a larger amount of rotations compared to WT MDCK cells. This suggests that larger adherence between cells either increases the coherence length or/and initiation is facilitated due to higher cell polarization rates μ and nu. Therefore, we can conjecture that the spontaneous length could be organ dependent and contribute to the key morphogenesis events.

### RhoA activity and its inverse relation with velocity

We report live activity of RhoA integrated over an entire multicellular system. With coarse graining, we considered MDCK rings as a continuum material where RhoA has its own dynamics and extracted FRET levels through time and space. Our results point to a simplification in their connection : RhoA and velocity are inversely related (along other work^40^) and this allows to consider only velocities for the model. This simplification would need to be further tested in other situations but it opens an interesting framework with a simple relation between RhoA activity and cell velocity. In future studies, RhoA and other Rho GTPases could be modulated through space and time using photoactivable tools to further understand its implication in collective coherent motion^41,42^. This will allow to control cell polarity and cell velocity and in turn characterize the coupling between Rho signaling and cell mechanics in specific manner with identification of new mechanosensory proteins.

### Implication of coherent motions *in vivo*

Our velocities found during coherent motion were close to the ones found during *Drosophila* egg chamber development ^10,11^. In this context, follicle cells collectively rotate along the chamber and the presence of lamellipodia is reported^10^. This could potentially mean that the phenomena described in this work on 2D rings, driven by cell “cryptic” lamellipodia, act also during these *in vivo* processes. Also, coherent rotations have been reported in various morphogenetic events *in vivo* in mammary acini, in egg chamber and in testis rotation in *Drosophila* ^10–13,43^. We propose that these phenomena could be generic and experienced by any growing tissues as suggested also by a recent theoretical study ^44^. If right, spontaneous rotation could occur when coherence lengths of each epithelial layer would be similar to dimensions of tissues. It would be interesting to compare coherence lengths across model systems to evaluate whether this generic property of living matter is physiological and is important for optimal development.

## Supporting information

Annex Math

## Acknowledgments

We thank J. Van Unen and M. Inamdar for discussions and feedbacks. We also thank A. Honigmann for kindly sharing the ZO1-GFP MDCK cell line, the Imaging Platform of IGBMC, and the Riveline Lab. for help and discussions. S.L.V. is supported by the University of Strasbourg and by la Fondation pour la Recherche Médicale. D.R., M.S. and L.N. acknowledge supports from Idex Unistra and from the Cell Physics Master at the University of Strasbourg. O.P. and D.R. thank funding from SNF Sinergia. This study has been also supported by a French state fund through the Agence Nationale de la Recherche under the frame program Investissements d’avenir labeled ANR-10-IDEX-0002-02.

## MATERIAL AND METHODS

### Micro-contact printing

Ring motifs are patterned on glass coverslips using micro-contact printing. Polydimethylsiloxane (PDMS; 1:10 w/w cross-linker:pre-polymer) (Sylgard 184 kit, Dow Corning, cat. DC184-1.1). Stamps are produced with UV-photolithography. Stamps are rinsed with 70% ethanol and then rendered hydrophilic by exposure to oxygen plasma (30s) (Diener Electronic, cat. ZeptoB). They are next incubated with a 10 μg/mL rhodamine-labelled fibronectin solution (Cytoskeleton, cat. FNR01-A) for 1h and then dried at room temperature for about 5min. In the meantime, glass coverslips are functionalized by vapour phase for 1h with 3-(mercapto)propyltrimethoxysilane (FluroChem). After incubation of both the coverslip and PDMS, the stamp is deposited onto the glass slide and kept in contact during 30min. To ensure a proper transfer between the two surfaces, a 50g weight is placed on the top of the stamp. Finally, the stamp is removed and the printed coverslip is washed with PBS 1X and stored in Milli-Q water. Non-printed areas are passivated with 0.1 mg/mL PLL-g-PEG (in 1mM HEPES pH 7.4, SuSoS AG, cat. SZ33-15) for 20min at room temperature. This approach consists in setting differential coating on the surface allowing to plate cells and select cells adhering to the fibronectin printed ring.

### Cell Culture

MDCK cells stably transfected with E-cadherin-DsRED are cultured in low glucose Dulbecco’s Modified Eagle’s (DMEM) with 10% Foetal Bovine Serum (FBS) and 1% Penicillin-Streptomycin. Cells are kept subconfluent (at ∼ 70%). Prior experiments, cells are detached with Trypsin-0.25% EDTA (Fisher Scientific, cat. 11570626) and centrifuged at 500rpm during 3min. The pellet is then re-suspended in DMEM 1% FBS. This low serum concentration will further impede attachment on the passivated surface during the first hours following seeding (differential adhesion). After 1h incubation, the printed surface is carefully washed out to remove cells from the non-printed area. After washout, DMEM is replaced by L-15 Leibovitz Medium (Fisher Scientific, cat. 11540556) supplemented with 1% FBS and 1% Penicillin-Streptomycin and the sample is incubated 1h under the microscope at 37°C before acquisition. This last incubation allows cells to spread on patterns and ensures that rings are closed at the onset of the experiment.

For laser ablation experiments, MDCK stably transfected with GFP-Myosin Regulatory Light Chain (MRLC) were used. Tight junctions were imaged with a MDCK cell line genetically engineered with CRISPR-Cas9 to express ZO1-GFP ^45^. We generated a MDCK cell line stably expressing the RhoA FRET biosensor using Piggy-Bac engineering. Briefly, cells were transfected with the biosensor plasmid together with a transposase. After integration into the genome, fluorescent cells were sorted with flow cytometry and kept under antibiotics pressure for 3 weeks.

### Transfection

Plasmids for mosaic experiments such as Actin-GFP or Caldesmon-GFP were incubated with cells using Lipofectamin (Invitrogen, cat. 11668030). One day prior experiment, cells at 50% confluency in a 6-well plate were incubated for 4-6h with 1μg of DNA and 10μL of lipofectamin mixed in Opti-MEM medium. After incubation, wells are washed out and the medium is replaced by fresh DMEM 10% FBS.

### Drug experiments

Cells were incubated with CK666 50μM (Sigma-Aldrich, cat. SML0006) during the whole course of the experiment (12 hours).

For RhoA inhibition, cells were incubated 2 hours with C3-transferase (Cytoskeleton Inc., cat. CT03-A) at a concentration of 1μg/mL. Culture medium was washed out before the start of the experiment.

### Standard time-lapse microscopy

Phase contrast images were acquired through a 10x objective (NA = 0.25) with a frame rate of 1 image every 10min. Patterns were systematically checked prior the time-lapse start with a standard epifluorescence lamp (FluoArc Hg Lamp) coupled with a rhodamine fluorescence filter. The very same set-up was used to acquire cells during mosaic experiments (actin-GFP and caldesmon-GFP). Tight junctions (ZO1-GFP) were imaged with a confocal microscope (Leica SP8 inverted) and an autofocus system. All z-planes were taken and finally projected onto one single plane.

### Biosensor FRET activity

Images were acquired with a confocal microscope (Leica SP8 inverted) with the pinhole opened to the maximum in order to increase slice thickness and the signal to noise ratio for the analysis. We used two Photo Multiplier Tubes (PMTs) as detectors. Bandwidths were chosen as follow: 472nm-512nm for mTFP1 (CFP analog, donor) and 523nm-544nm for Venus (YFP analog, acceptor). mTFP1 was excited by an argon laser at 468nm with 15% of power and emission for mTFP1 (donor) and Venus (FRET) were simultaneously acquired by the two PMTs. Images were then corrected and processed with Biosensor Package from Danuser Lab (available on the following link: https://github.com/DanuserLab/Biosensor). Because it is a unimolecular biosensor, no bleedthrough correction was done. Prior analysis, histograms were manually resliced in order to remove high and close-to-zero boundary artefacts.

To check whether the FRET sensor was reliable, we performed additional tests: incubation with 1.0 μg/ml C3-transferase (RhoA inhibitor, Cytoskeleton, cat. CT04) and with 1.0 μg/ml of RhoA activator (Cytoskeleton, cat. CN03). Drugs led to decrease and increase of activities respectively (Fig. S4a-b). We also used a mutant sensor – F39A, unable to interact with the RBD domain ^18^ – with the same acquisition parameters detailed above. This was associated with a significant decrease in the FRET signal (Fig. S4c).

### Laser ablation

Laser ablations on acto-myosin cables were done using the FRAP module from a Leica SP-5 inverted microscope. The sample was imaged and ablated through a 40x oil objective (NA = 1.25) and an 800nm infrared laser (80 MHz, pulsed).

### Velocity fields and coherence length

Velocity fields were generated using PIVlab 1.4 and performing cross-correlations. Windows of about a cell size, adjusted to the magnification (32×32 or 64×64 in PIVlab) were taken. No smoothing was performed on the velocity fields.

Coherence lengths *ξ* were extracted from velocity fields with a custom-made Matlab code by computing the spatial correlation function of the tangential velocity *v*_*θ*_:

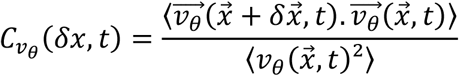

Exponential decays are generated from the computations of 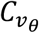 and the intersections between the tangents at origin and x-axis give *ξ*. Because 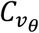 was not always converging to 0, we made linear fits on the 3 first points of 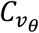 in order to extract the tangent at origin.

We selected *ξ*_*max*_ to capture the largest length accessible to the cells. To extract this maximal coherence length *ξ*_*max*_ presented in Figure 1c and Fig. S1a, velocity fields were averaged over intervals of 10 frames (100min) and the time interval leading to the largest correlation function was considered. Stated differently, we isolated the time window during which the coherent motion was optimal and extracted the associated length over experimental conditions.

### Image processing

Nuclei in RhoA ratiometric maps were removed using Ilastik and phase contrast images were processed with Fiji.

Cell polarity was defined by its lamellipodium: the axis nucleus-lamellipodium sets direction and orientation for cell polarity. Since many factors could contribute to define physical polarity and because lamellipodia are not always observable, we decided to track ZO1 marked cells – not in their anisotropy of signal – but for the well-defined shapes. In majority of cases, cells could be associated to isocele triangles. Therefore, cell polarity was given by the long axis of the triangle while the sense was defined by the triangle base. We confirmed that this polarity was consistent with the polarity defined by lamellipodia in the large majority of cases (Fig. S2a-b). Segmentation and extraction of single cell parameters in ZO1 experiments were done using Tissue Analyzer ^46^ in Fiji.

### Model

We designed a mechanistic-based minimal “Vicsek—type model” in which cells are treated as particles with polarity **p** and velocity **v**, initially placed on a ring with two layers of cells matching the cell density and number tested in experiments. Their interaction rules are determined by alignment of polarities and by polarity relaxation to the direction of motion and by random fluctuations of polarities (see Annex Math for details). Velocities are also projected on the directions of polarities to the limit of volume exclusion. The inner and outer supra-cellular acto-myosin cables are modeled as segments connecting boundary cells with two physical elastic properties, stretching and bending energies. In addition to the force exerted by the cables on the cells and its velocities, cables also act on cell polarity. The values of parameters are extracted from measurements and from estimates based on references (see Annex Math and Table 1).

We show (see Fig. 5e) that with this passive treatment of cables and actual experimental values, we do not recapitulate coherent motion. Based on the Rho activity at both cables supporting an active stress generation, we add in the model an active cable force with estimates of the associated experimental values. Simulations based on the theoretical model reveal that cells undergo coherent motion for small rings. Moreover, when we increase the ring diameter keeping the same values for parameters, we reproduce the experimental dynamics with no further adjustments. These results support the relevance of our model.

## Notes

### Competing Interest Statement

The authors have declared no competing interest.

### Summary of Updates

This version of the manuscript has been revised to update experiments and simulations.

