## Supplementary Material for "Spontaneous rotations in epithelia as an interplay between cell polarity and boundaries"

### Supplementary Materials

#### Coherence and fluidization

To undergo coherence, cells need to change neighbours through T1 transitions. We report these events in Fig. S2e, and we show that the most coherent rings exhibit the largest T1 rates even though the T1 rates were constant for a given experiment (Fig. S2f). This is consistent with the fact that the denser rings had the largest coherence (Fig. S2g). This constant T1 rates was also observed in non-coherent rings but their cumulative numbers were lower than coherent rings, which suggests that the coherent rings have a more fluid behaviour than non-coherent.

Altogether, cell density and cell divisions are involved in the emergence of collective coherent motion and this is summarized in the phase diagram shown in Figure S2h.

#### 2D versus 1D: ring thickness, neighbour exchanges and radial motion

As detailed above with T1 transitions, neighbour exchanges impact the onset of coherence and these transitions occur because the thickness of the ring pattern (30µm) allows to have locally two rows of cells. Therefore, the presence of these phenomena leads us to consider the ring as a 2D structures even though only tangential motions are extensively analysed as it is the essential readout for coherent motion.

Because the local curvature is high in coherent rings (diameter = 80µm) and because of the contractile cables (see below for the ellipse case), cells tend to have a radial component to their motion that closes the ring. This generates a disk-like structure. We believe that the dynamics emerging in this new configuration is distinct from the one in the ring configuration. Therefore, we excluded this state for the analysis and the comparison with our theoretical model.

#### Effects of cables and curvature

We think that local curvature is also an important component in preventing cells to extend outwards, mainly through the passive component of the cable. These observations and hypothesis could potentially explain the radial extensions in finger-like structures for large rings (1000µm): the local curvature is extremely low compared to the cell size. Therefore, extension in and out are equivalent and no direction is favored through the constraint generated by the cables.
The effect of cables is further tested with experiments with a modified initial geometry. We plated cells on fibronectin micro-contact printed ellipses. Dynamics was distinct around the cell layer : the closure happens faster at edges where the local curvature is larger (Fig. S3B).This supports again a mechanical role of the cable.

#### Radial component for motion

This study focuses on the emergence of coherent motions in 2D cellular rings. Therefore, the main readout for coherence is the tangential component of the velocity field (which is used to extract correlation functions and coherence length). We mainly considered this parameter and we tested different conditions and generic interaction rules to understand what dictates its value. Nevertheless, we are aware that cells also exhibit radial motions since migration is not confined between physical walls. This component seems to be mainly controlled by the characteristics of the cables and especially by the bending and the stretching moduli, as discussed above: we observe in larger rings (1mm) the emergence of finger-like structure driven by leader cells. Appearance of leader cells also occurs when acto-myosin cables are altered by incubation with caldesmon. Contribution of radial motion will be interesting to characterize in future experiments.

#### Interplay between RhoA level and coherence

We looked for potential links between the onset of coherence and FRET levels. We saw that rings with higher FRET at initial time $t_0$ could not reach coherence (Fig. 4f). To modulate the level of Rho, we used C3-transferase, a RhoA inhibitor, to decrease the global RhoA level. Rings with C3-transferase did not undergo rotation and their coherence was low ($\frac{\xi}{p}$ $0.4$). By comparing all rings, we could extract three populations (Fig. S4i, left) based on the $\frac{\xi}{p}$ levels together with the RhoA activity. The resulting binned curve (Fig. S4i, right) shows an optimal $\frac{\xi}{p}$ for a medium RhoA activity. These results allow to draw a non-linear relationship RhoA – coherence that could be potentially explained by frictions arguments as suggested in previous works for cell velocity and adhesion (Palecek *et al*, 1997).

#### Polarity strength and lamellipodia extension

In this study, we consider for the sake of simplicity that every lamellipodium within the ring contributes equally. In future work, it would be interesting to convert the extension of the lamellipodia in a polarity value to test their contributions.

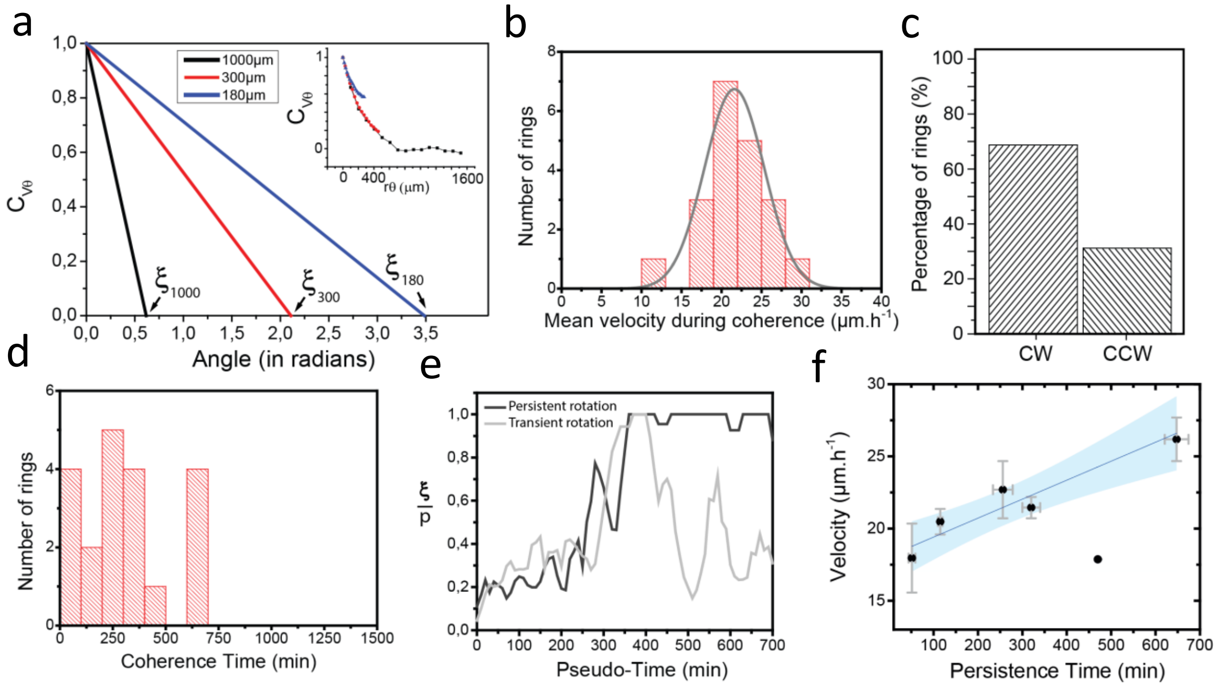


**Figure S1. Different populations of coherent rings.** (a) Tangents at origin from the correlation functions of the tangential velocity gives $\xi_{max}$ for each configuration. Insert shows the original plot of the correlation as a function of curvilinear length. (b) Distribution of ring velocities during coherence. (c) Clockwise (CW) orientation is favored during rotation compared to the counter-clockwise (CCW) orientation. (d) Distribution of coherence durations, assessed by Xsi/p going from 1 down to lower than 0.9. (e) Two ring populations appear after the onset of coherence: (i) rings undergoing persistent rotation and (ii) rings exhibiting transient rotation with a fast decay of the coherence index. (f) Coupling between ring velocity and persistence of coherent motion. Data are binned with respect to time. Timers in hh:mm. Scale bars = 50µm.

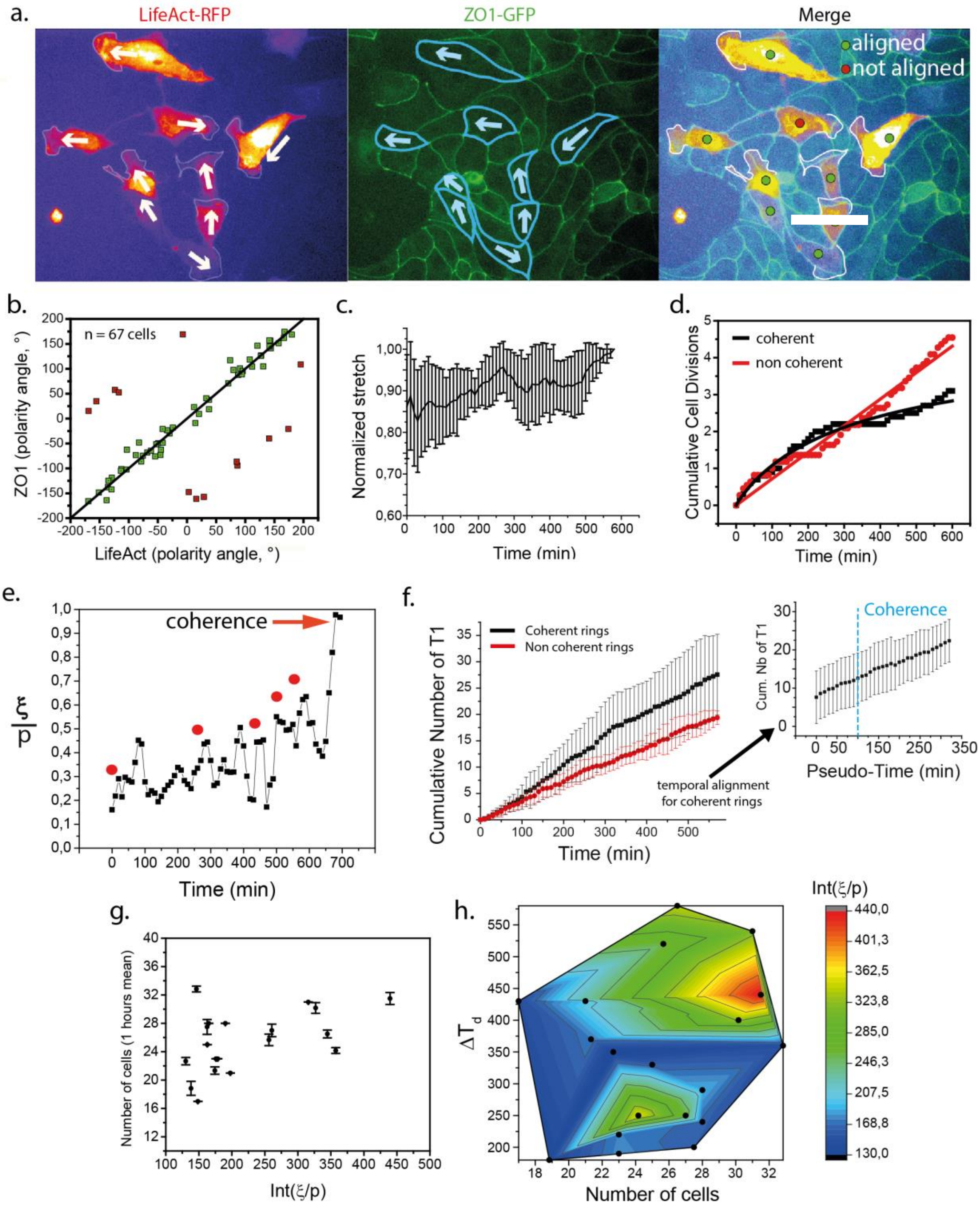


**Figure S2A. Extraction of single cell parameters within the ring.** (a) Epithelium of MDCK cells expressing ZO1-Ng transfected with LifeAct-RFP. Actin visualization allows to identify lamellipodia and extract cell polarity. Scale bar = 20µm. Cell polarity given by the lamellipodium and determined by ZO1 distribution are compared in (b). Green = correlated, red = non-correlated. (c) Normalized stretch as a function of time for non-coherent rings. (d) Complete distributions of cumulative cell divisions shown in Fig. 2. (e) Impact of cell divisions on coherence initiation. Each red dot corresponds to a cell division within the ring. (f) Cumulative number of T1 transitions in coherent and non-coherent rings. The inset shows the variation of T1 transitions with respect to the coherence initiation. (g) Number of cells as a function of the integer of $\frac{\xi}{p}$ as a readout for coherence during the experiment. (h) Phase diagram including three parameters: $\Delta T_d$ the maximal time between two divisions, the number of cells and the integer of $\frac{\xi}{p}$.

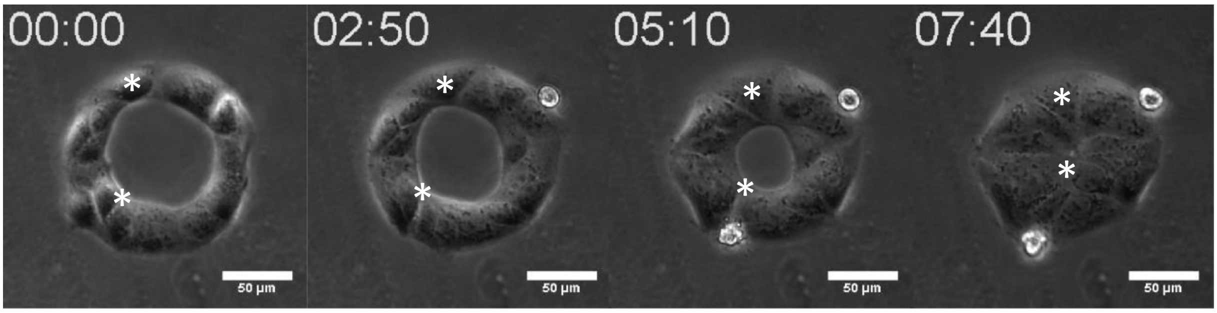


**Figure S2B. Tug-of-war inhibition.** Cells were incubated with Arp2/3 inhibitor CK666 and this prevented coherence initiation and promoted radial closure in a rosette-like manner. Stars show two points in the ring and this indicates the absence of rotation. Time in hh:mm. Scale bar = 50µm.

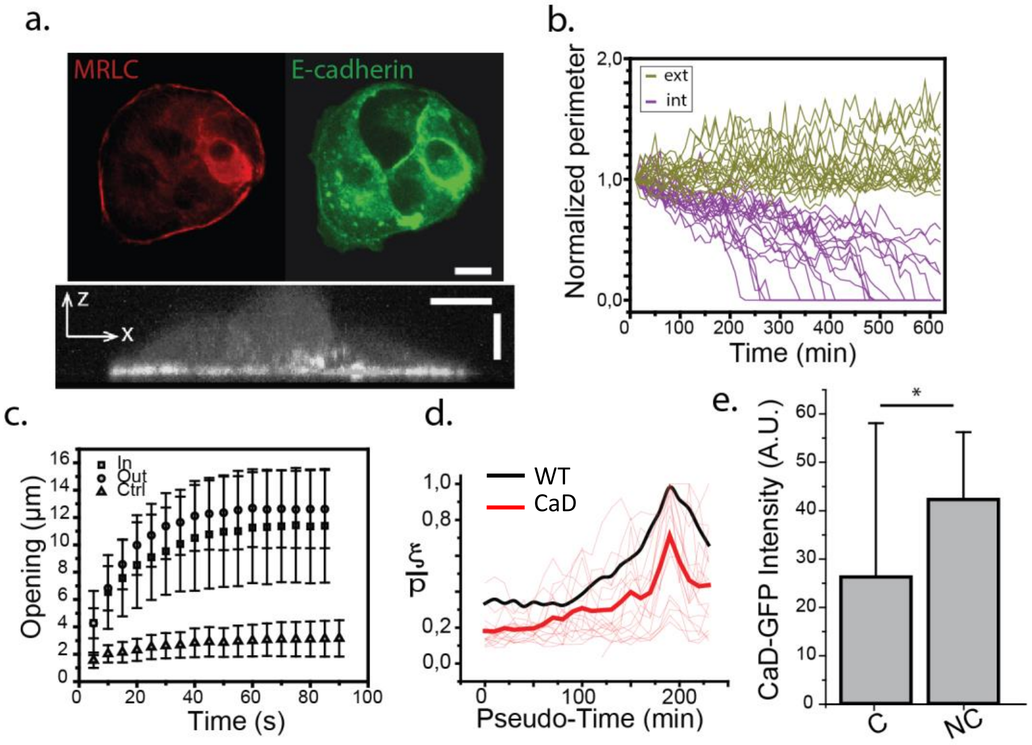


**Figure S3A. Acto-myosin cables assembly and dynamics.** (a) Acto-myosin cable at the boundary of WT MDCK colony on a glass surface. Scale bars = 5µm. (b) Dynamics of inner and outer boundaries of 80µm rings (c) Opening dynamics with laser ablation of acto-myosin cables and cell-cell junction (control-ctrl). (d) Coherence dynamics for CaD rings and WT rings. (e) Fluorescence intensity of CaD transfected cells within the ring for coherent (C) and non-coherent rings (NC). Coherent rings correspond to lower CaD expressions.

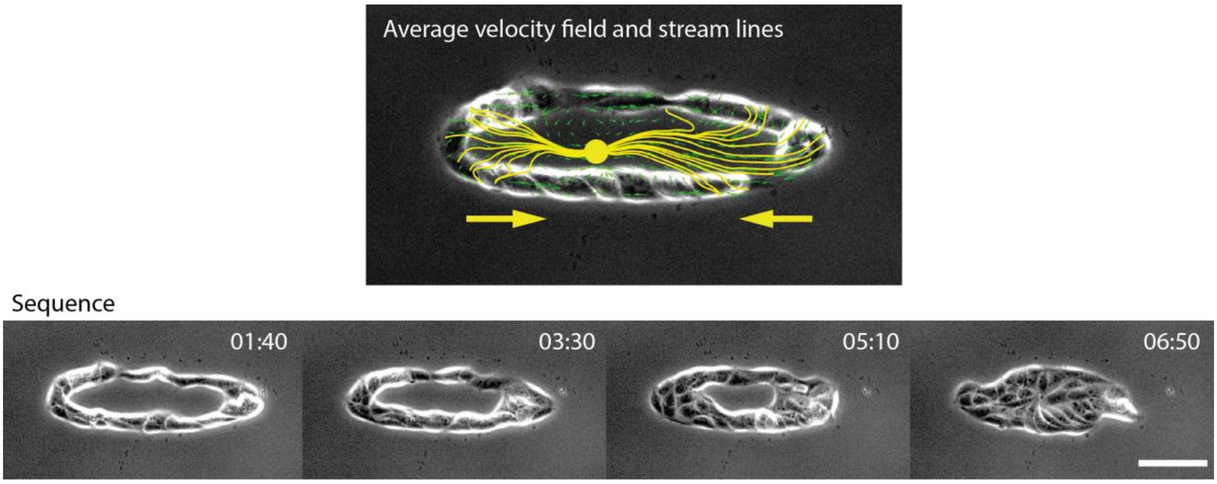


**Figure S3B. Local curvature and closure.** MDCK ellipses exhibit faster closure at edges where the local curvature is the highest. Stream lines are shown in yellow, average PIV in green. Time in hh:mm. Scale bar = 100µm.

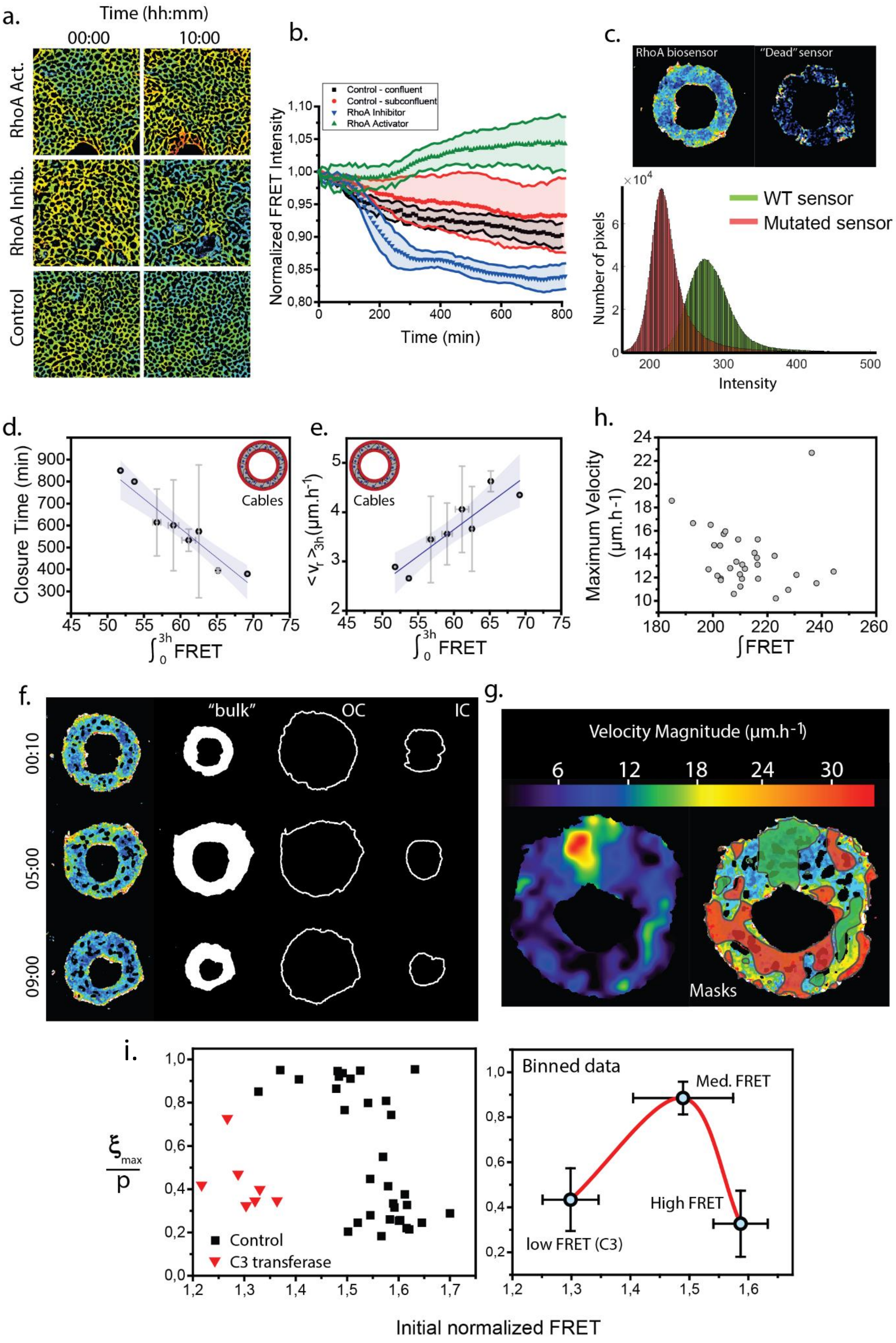
a.
Time (hh:mm)
00:00
10:00
RhoA Act.
RhoA Inhib.
Control
b.
Control - confluent
Control - subconfluent
RhoA Inhibitor
RhoA Activator
Normalized FRET Intensity
Time (min)
c.
RhoA biosensor
"Dead" sensor
WT sensor
Mutated sensor
Number of pixels
Intensity
d.
Closure Time (min)
Cables
e.
Cables
h.
Maximum Velocity (µm.h-1)
∫FRET
f.
"bulk"
OC
IC
00:10
05:00
09:00
g.
Velocity Magnitude (µm.h-1)
Masks
i.
Control
C3 transferase
Binned data
Med. FRET
High FRET
low FRET (C3)
Initial normalized FRET

**Figure S4. RhoA activity in rings.** (a) MDCK epithelia treated with different RhoA regulating drugs together with the FRET signal quantification over time in (b). (c) FRET intensity acquired with a mutated sensor (described in ref [18]) shows a decrease in intensity. The set of experiments in panels (a-c) confirms the efficacy of our RhoA sensor and the proper images acquisition and analysis protocols. (d) Average FRET level at the cables, integrated over the initial time points (3h) correlated with the closure time. High RhoA activity at boundaries is associated with a faster ring closure. This read-out can be assessed by (c) where the average radial velocity is compared to the initial average FRET level. (f) Segmentation of the multicellular rings. Two regions are distinguished for analysis: the acto-myosin cables at the two interfaces and the cellular "bulk" within the two cables. (g) Velocity regions defined as low ($< 5 \mu m.h^{-1}$) and high ($> 15 \mu m.h^{-1}$) associated with their masked FRET map (green mask = high velocity, red mask = low velocity). (h) Full data of the FRET analysis shown in Fig. 4e. (i) Coherence index as a function of the initial normalized FRET.

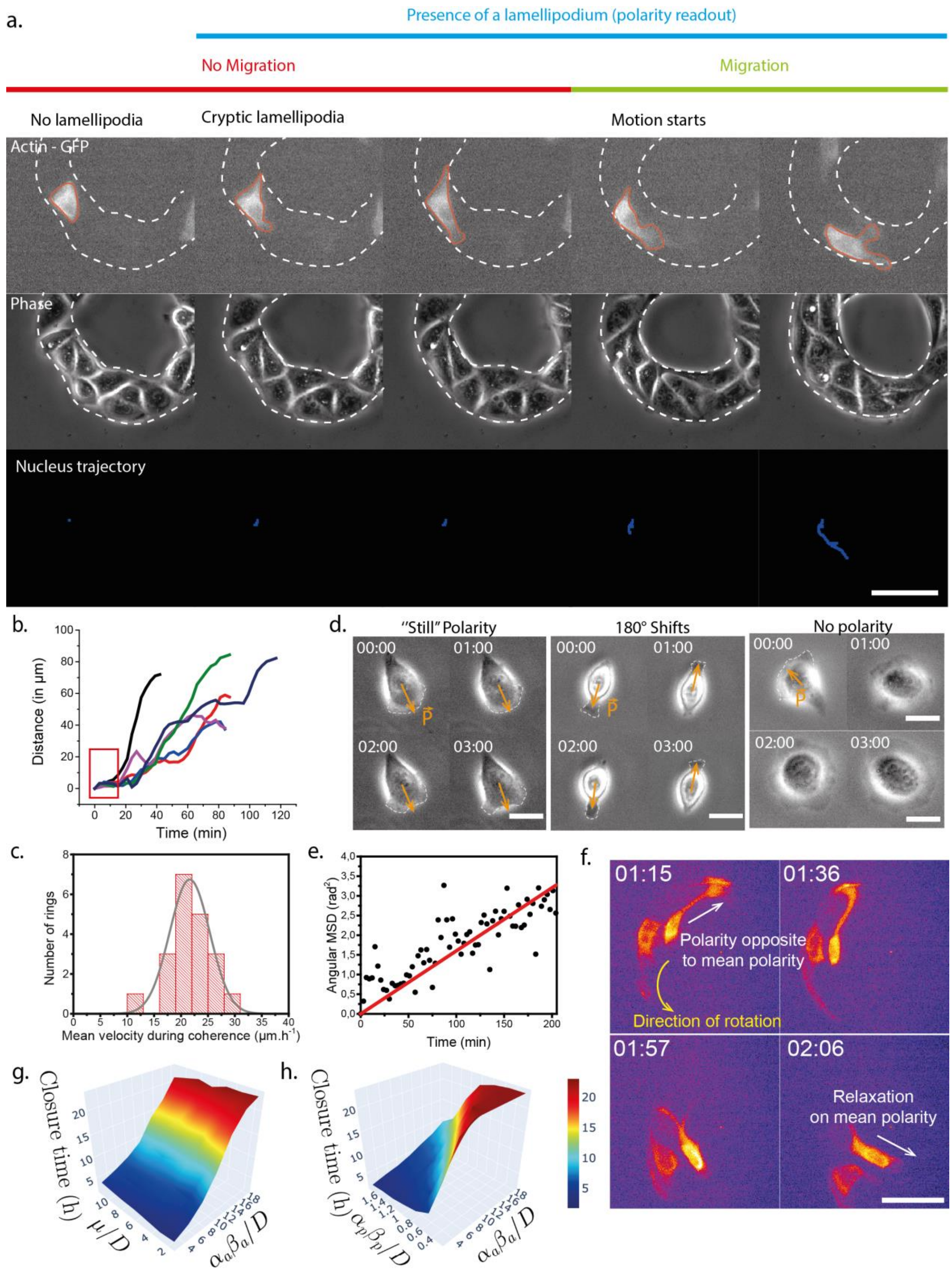


**Figure S5. Model characterization.** (a) Tracking of cells before lamellipodium growth within the ring. Polarity set by the lamellipodium precedes cell motion as assessed by nuclei displacement shown in (a) and (b). Scale bar = 50µm.(c) Distributions of mean ring velocities during coherence. The Gaussian fit sets the values of maximal cell velocities $c$ in the model. (d) Single cell polarity diffusion and the associated MSD and diffusion coefficient $D$ shown in (e). Scale bar = 15µm. (f) Experimental definition of cell relaxation rate $\mu$. This quantity reflects the velocity at which a given cell polarized in the direction opposite to the mean polarity $\overline{P_k}$ relaxes to $\overline{P_k}$. Scale bar = 20µm. (g-h) *Phase diagrams relative to the ring closure time (in hours) with respect to the dimensionless parameters associated with the model (active cable force - active alignment, active cable force – passive cable force). Index p refers to the passive component of the cable while a corresponds to the active one.* On each plot, each point of the surface corresponds to the average value of 50 simulations.

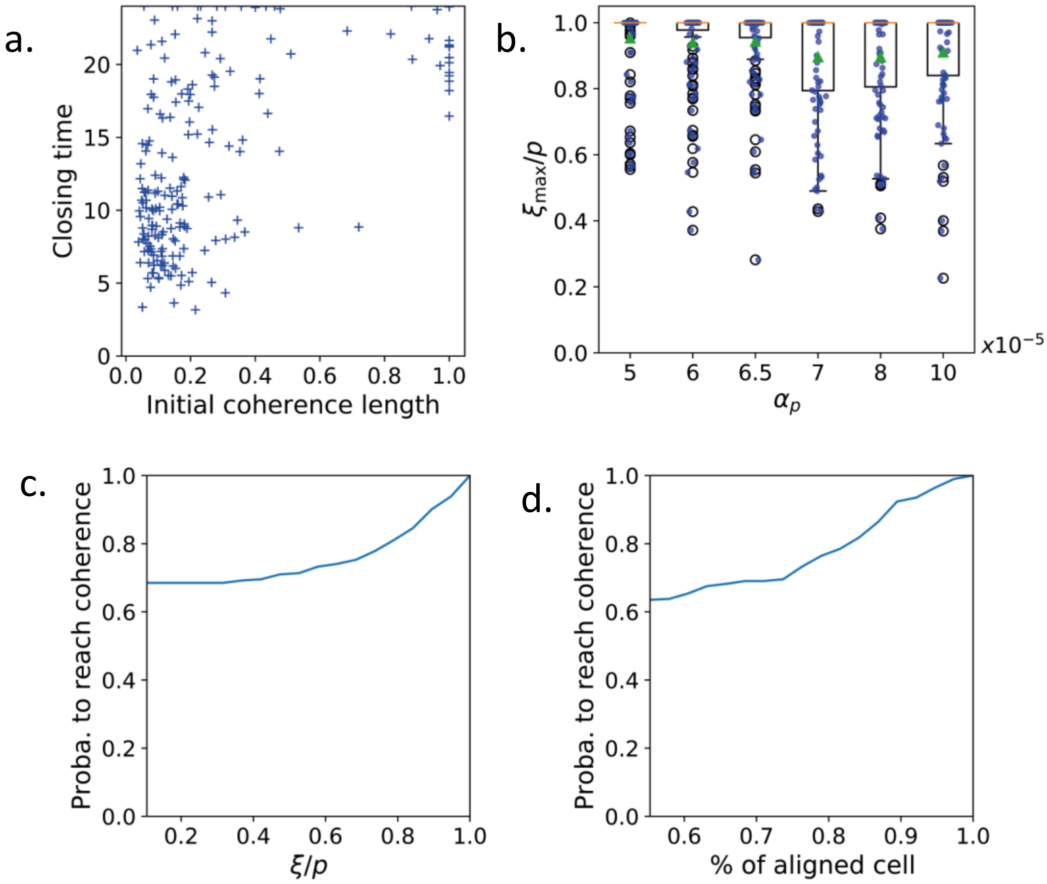


**Figure S6. Additional analysis from the model.** (a) Closing time as a function of the initial coherence length. (b) Coherence index as a function of the cable passive term value. (c) Probability to reach coherence based on the global coherence and (d) as a function of the percentage of aligned cell within the ring.

| Parameter | Symbol | Value | Units |
|---|---|---|---|
| Particle radius | $r$ | 7.5 | µm |
| Velocity | $c$ | 21.6 | $\mu m.h^{-1}$ |
| Angular diffusion coefficient | $D$ | 0.96 | $rad^2.h^{-1}$ |
| Stretching modulus | $k_{st}$ | $5.10^3$ | $pN.\mu m^{-1}$ |
| Bending modulus | $k_b$ | $10^3$ | pN.µm |
| Relaxation rate on $\overline{P_k}$ | $\mu$ | 6.2 | $rad.h^{-1}$ |
| Relaxation rate on $\frac{V_k}{\lVert V_k \rVert}$ | $\nu$ | 6.2 | $rad.h^{-1}$ |
| Active force | $f$ | 10 | nN |

**Table 1. Single cell parameters used in the theoretical model.** See also Sec. 2.2 from Annex Math.
