## Supplementary material for "Spontaneous rotations in epithelia as an interplay between cell polarity and boundaries": Annex Math

### A mathematical model for self-confined cells dynamics (Annex Math)

In this part, we provide details about the particle model we designed for the numerical simulations of cell aggregates. It is a Vicsek-type model in interaction with self-generated surrounding acto-myosin cables. The derivation of the model is guided by experiments.

#### 1 Continous model: derivation and scaling analysis

##### 1.1 Geometrical description of cells and cables

In a first approximation, cells are considered as hard-disks of radius $r > 0$ and their deformation is neglected. Initially, we consider $N$ such particles that are distributed in a two-dimensional ring domain: $\Omega = \{(r, \theta) \in [R_{\min}, R_{\max}] \times [0, 2\pi[\}$. This initial configuration is depicted in Fig. A1a.

The $k$-th cell dynamics is described by three vectors: its position $X_k(t) \in \mathbb{R}^2$, its velocity $V_k(t) \in \mathbb{R}^2$ and its polarity $P_k(t) \in \mathbb{S}^1$, where $\mathbb{S}^1$ denotes the set of unit vectors $\mathbb{S}^1 = \{d \in \mathbb{R}^2, \|d\| = 1\}$ and $\|\cdot\|$ is the Euclidean norm, see Fig. A1b for a zoom. Here, polarity is defined by the direction of lamellipodia. Since we consider a minimal model, this polarity has no varying magnitude. We denote by $\mathbf{X}(t) = (X_k(t))_{k=1,\ldots,N} \in (\mathbb{R}^2)^N$ the vector of all positions, by $\mathbf{V}(t) = (V_k(t))_{k=1,\ldots,N} \in (\mathbb{R}^2)^N$ the vector of all velocities and by $\mathbf{P}(t) = (P_k(t))_{k=1,\ldots,N} \in (\mathbb{R}^2)^N$ the vector of all polarities, respectively.

As observed in experiments, acto-myosin cables appear at boundary of multi-cellular rings. These cables are intracellular and form a continuous line that encloses cells and acts as an active physical boundary. From our modeling point of view, these cables are defined as piece-wise affine curves linking the cells located on the inner and the outer boundaries, see Fig. A1 for a schematic. We thus need to identify frontier cells within the ring domain. The identification algorithm is based on the hypothesis that cells remain in a disk-like or a ring-like configuration over time and that the two curves can be described as graphs with respect to a polar angle. We consider polar coordinates with respect to the barycentre of cell positions and at each discretized polar angle is thus associated the cell with minimal polar radius and the one with maximal polar radius.

We denote by $N_{\text{in}}$ (resp. $N_{\text{out}}$) the number of cells belonging to the inner (resp. outer) boundary at time $t > 0$ and by $\pi_{in}$ and $\pi_{out}$ two sets of size $N_{\text{in}}$ and $N_{\text{out}}$ that provide the indexes of the boundary cells ordered by increasing polar angle (still in the reference of the barycentre). Then the cables are defined as the polygonal deformable curves:

$$\mathcal{P}_{\text{in}} = X_{\pi_{\text{in}}(1)} X_{\pi_{\text{in}}(2)} \cdots X_{\pi_{\text{in}}(N_{\text{in}})}, \tag{1}$$

$$\mathcal{P}_{\text{out}} = X_{\pi_{\text{out}}(1)} X_{\pi_{\text{out}}(2)} \cdots\cdots X_{\pi_{\text{out}}(N_{\text{out}})}. \tag{2}$$

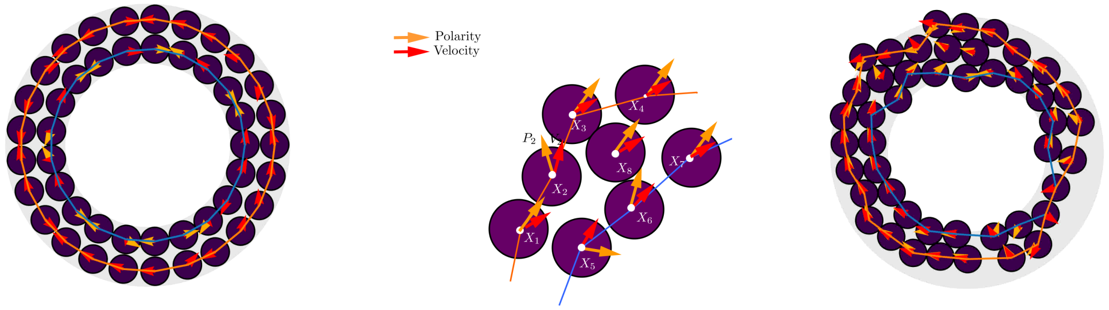


(a) Initial configuration.

(b) Notations: positions ($X_k$), velocities ($V_k$), polarities ($P_k$).

(c) Typical configuration close to the onset of the numerical simulation after t=0.

Figure A1: Geometrical configuration. Velocity vectors (red), polarity vectors (orange), inner cable (blue), outer cable (orange).

Note that vectors $\pi_{in}$, $\pi_{out}$ and consequently these polygonal curves change constantly in time as cells can move from the boundary to the core of the ring and vice-versa, as it can be observed in Fig. A1c. However, we do not explicitly write the dependency with respect to time. More details about the biophysical mechanisms underlying the deformation of the cables will be provided in the next section.

#### 1.2 Mathematical modeling of cells and cables dynamics

**Positions and velocities dynamics.** The particle velocities $V_k$ are governed by the direction of the polarity $P_k$ up to a contact force that prevents from overlapping. The relation between velocity and polarity is supported by experiments, see Figure S5. The fundamental laws of the dynamics then write:

$$\frac{dX_k}{dt} = V_k,$$
$$\varepsilon m \frac{dV_k}{dt} = \frac{1}{\gamma}(cP_k - V_k) + (F^{\text{cable}}_{\text{passive}})_k + (F_{\text{contact}})_k,$$

where $c > 0$ is the maximal velocity amplitude, $\gamma$ is the inverse friction coefficient and $\varepsilon > 0$ is the ratio between inertia and viscous forces. On the right-hand side of the velocity equation, the first force is directed along the polarity $P_k$ and corresponds to an active force, the second term -$\frac{1}{\gamma}$ $V_k$ is the friction force resulting from the cell environment, the third one is associated to the inner and outer cables (defined below in Eq.(9)) and the last one stands for contact forces due to the volume exclusion constraint. In the overdamped limit $\varepsilon \to 0$, we have :

$$V_k = cP_k + \gamma\,(F^{\text{cable}}_{\text{passive}})_k + \gamma\,(F_{\text{contact}})_k.$$

Therefore, the contact force can be interpreted as the Lagrange multiplier associated to the volume exclusion constraint. Specifically, by denoting $\mathbf{F}_{\text{contact}} = ((F_{\text{contact}})_k)_{k=1,\dots,N} \in (\mathbb{R}^2)^N$ the vector of all forces, the term $\gamma \mathbf{F}_{\text{contact}}$ equals the forces such that $\mathbf{V}$ are the velocities closest to $c\mathbf{P}$ that do not lead to overlapping. Following the framework studied in [16, 17], this can be reformulated by writing that the desired velocity $c\mathbf{P}$ is projected onto a nearest admissible velocity that prevents the overlap. Thus, by gathering together equations for all cells, for $k = 1, \ldots, N$, the previous system becomes:

$$\frac{d\mathbf{X}}{dt} = \mathbf{V}, \tag{3}$$

$$\mathbf{V} = \mathbf{Proj}_{\mathcal{C}_{\mathbf{X}}}\Big(c\mathbf{P}\Big), \tag{4}$$

where $\mathbf{Proj}_{\mathcal{C}_{\mathbf{X}}}$ denotes the projection operator onto the set of admissible velocities defined by:

$$\mathcal{C}_{\mathbf{X}} = \left\{\mathbf{V} \in (\mathbb{R}^2)^N | \quad \forall i < j, \quad D_{i,j}(\mathbf{X}) = 0 \quad \Rightarrow \quad \nabla D_{i,j}(\mathbf{X}) \cdot \mathbf{V} \geqslant 0\right\},$$

where $D_{i,j}(\mathbf{X}) = \|X_i - X_j\| - 2r$ corresponds to the distance between the $i$-th and $j$-th particles of radius $r$. Note that it is non-local operator as this set depends not only on the positions and velocities of each particle, but accounts for the neighbours within the cell ring. Hence in this model, the velocity instantaneously takes the direction of cell polarity unless contact forces compete. Velocities magnitudes depend on jamming: cells move with maximal velocity speed $c$ when cell alignment is optimal.

**Polarities dynamics.** Several mechanisms are incorporated to account for the polarity dynamics. First, the polarities relax to the velocity unit direction $\frac{V_k}{\|V_k\|}$, at angular rate $\nu > 0$. In addition, they have active interactions with the neighboring polarities and with the acto-myosin cables. More precisely, the polarity of the $k$-th cell relaxes to the mean polarity $\bar{P}_k$ of the neighbouring particles with a angular rate $\mu > 0$,

$$\bar{P}_k = \frac{\sum_{j, \|X_j - X_k\| \leqslant 4r} P_j}{\left|\sum_{j, \|X_j - X_k\| \leqslant 4r} P_j\right|} \tag{5}$$

and reacts to the passive forces exerted by the cables $(F^{\text{cable}}_{\text{passive}})_k$, with an angular rate $\alpha_p > 0$ where the subscript $p$ stands for passive. Here we assume that these interactions can occur only with cells that are at a distance smaller than $4r$ (not more than two cell diameters apart). Moreover, the cables are composed of actin and myosin which behave as an active gel, intrinsically out-of-equilibrium [23]. In addition, FRET experiments showed high levels of RhoA at the boundaries suggesting a high activity at acto-myosin cables (see Fig. 4). Therefore, considering acto-myosin cables as only passive elastic materials would not be consistent with their active properties. The polarity also reacts to an active force $(F^{\text{cable}}_{\text{active}})_k$ with an angular rate $\alpha_a > 0$ where the subscript $a$ stands for active. Without this active force, we show that the system does not undergo coherent rotating motions in the physical parameters range (see Fig. 5-e). Taking these assumptions together, we obtain the following polarity equation:

$$\begin{aligned} dP_k = \mathbf{Proj}_{P_k^\perp} \circ \Bigg( \nu \left( \frac{V_k}{\|V_k\|} - P_k \right) dt + \mu \left(\bar{P}_k - P_k\right) dt \\ + \alpha_p \,(F^{\text{cable}}_{\text{passive}})_k \, dt + \alpha_a \,(F^{\text{cable}}_{\text{active}})_k \, dt + \sqrt{2D}\,(dB_t)_k \Bigg). \end{aligned} \tag{6}$$

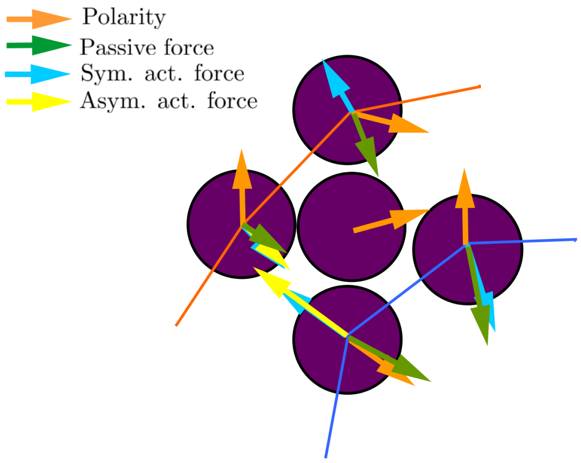


Figure A2: Forces acting on polarities (orange arrows). Inner cable in blue. Outer cable in orange.

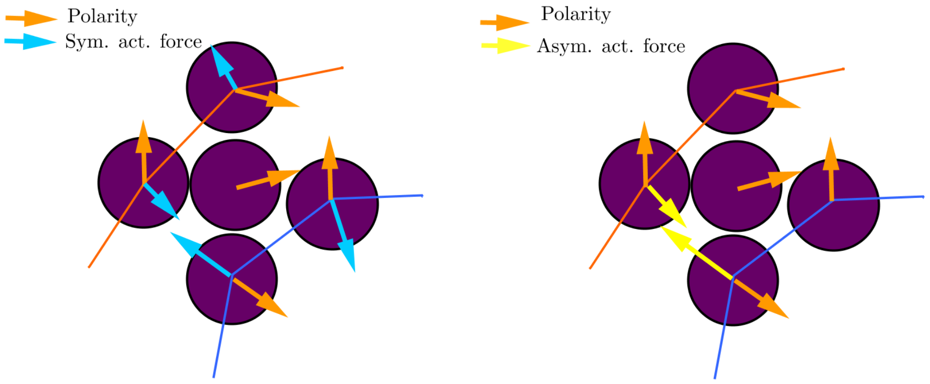


(a) Symmetric active force (blue arrows)

(b) Asymmetric active force (yellow arrows)

Figure A3: Active forces depending on polarity (orange arrows) and cables (piece-wise straight lines). Inner cable in blue. Outer cable in orange.

where the last term is a Gaussian white random noise with an angular diffusion coefficient $D > 0$ and the projection operator $\mathbf{Proj}_{P_k^\perp}$ ensures that the polarity remains of norm one over time by projecting the possible infinitesimal variation onto the orthogonal space to the vector $P_k$. This formulation corresponds to a Vicsek modeling approach, see for instance [10] for a review of such developments in the context of cell dynamics modeling.

**Acto-myosin cables dynamics.** We define in the next paragraph the contributions of the cables to the polarity dynamics by means of passive and active forces, which are schematically summarized in Fig. A2. Note that the arrows corresponding to different vectors are not represented to scale. We see that the four configurations (outward/inward polarity vectors versus inner/outer cable) lead to four different configurations of forces, detailed hereafter.

The acto-myosin cables forces are considered as ensembles of small springs connecting each particle at the outer $\mathcal{P}_{\text{out}}$ and inner $\mathcal{P}_{\text{in}}$ boundaries defined in Eqs. (1)-(2). Note that in a different context, such a description has been used to describe vesicles dynamics [11]. The energy of each cable is defined as:

$$E_{in/out}(\mathbf{X}) = \sum_{j=1}^{N_{in/out}} k_b(1-e_{\pi_{\text{in/out}}(j)} \cdot e_{\pi_{\text{in/out}}(j+1)}) + \sum_{j=1}^{N_{in/out}} k_{st} \left(\|X_{\pi_{\text{in/out}}(j+1)} - X_{\pi_{\text{in/out}}(j)}\| - 2r\right)^2, \tag{7}$$

with

$$e_{\pi_{\text{in/out}}(j)} = \frac{X_{\pi_{\text{in/out}}(j)} - X_{\pi_{\text{in/out}}(j-1)}}{\|X_{\pi_{\text{in/out}}(j)} - X_{\pi_{\text{in/out}}(j-1)}\|}, \tag{8}$$

the unit vector between two consecutive cells in the outer and inner cables, denoted hereafter $\mathcal{P}_{\text{out}}$ and $\mathcal{P}_{\text{in}}$, respectively. The first term corresponds to the bending energy of the springs, hereafter denoted $E^b_{in/out}(\mathbf{X})$, with bending modulus $k_b > 0$ and the second term corresponds to the stretching energy denoted $E^{st}_{in/out}(\mathbf{X})$ with stretching modulus $k_{st} > 0$. For the stretching part, the elongation is computed with respect to the contact case: the elongation is zero when the two cells are touching each other. Considering this energy, acto-myosin cables exert on any boundary particles a passive force given by:

$$(F^{\text{cable}}_{\text{passive}})_k = \begin{cases} -\nabla_{X_k} E_{in/out}(\mathbf{X}), & \text{if } k \in \pi_{in} \cup \pi_{out}, \\ 0, & \text{otherwise,} \end{cases} \tag{9}$$

and represented by a green arrow in Fig. A2. Note that this force is neglected in the velocity equation (4), but we assume that it has a significant contribution in the polarity dynamics to realign the cells tangentially to the cable. As already mentioned, the cables exert an additional active force on polarity. This active cytoskeletal force is decomposed into a symmetric and an asymmetric part as follows:

$$(F^{\text{cable}}_{\text{active}})_k = \begin{cases} (F^{\text{cable}}_{\text{active-sym}})_k + (F^{\text{cable}}_{\text{active-asym}})_k, & \text{if } k \in \pi_{in} \cup \pi_{out}, \\ 0, & \text{otherwise,} \end{cases} \tag{10}$$

represented by the blue and the yellow arrows in Fig. A2, respectively. The symmetry refers to the angle between the polarity and the cable and two contributions are incorporated, as described in detail in Fig. A3. Specifically, either (i) the polarity aligns to the cables whatever its polarity direction is and it leads to the symmetric force, see Fig. A3a:

$$(F^{\text{cable}}_{\text{active-sym}})_k = \beta_{\text{sym}} \left(\text{sgn}(t_k \cdot P_k)\, t_k - P_k\right), \tag{11}$$

where $t_k$ refers to the unit tangent vector to the cable, or (ii) the polarity aligns to the cables only when it is outward oriented with respect to the ring and it leads to the asymmetric force, see Fig. A3b:

$$(F^{\text{cable}}_{\text{active-asym}})_k = \begin{cases} \beta_{\text{asym}}\, n_k, & \text{if } (n_k \cdot P_k) > 0, \\ 0, & \text{otherwise,} \end{cases} \tag{12}$$

where $n_k$ refers to the unit inward normal vector to the cable. This force is thus always directed inwards with respect to the ring. We assume that each active force given in

Eqs. (11) and (12) has a constant order of magnitude, denoted $\beta_w$ and $\beta_n$, respectively. However the magnitude of the symmetric force depends on the alignment to the cable, while the asymmetric force has a constant magnitude when the polarity points outward the ring. The tangent vector $t_k$ and the inward tangent vector $n_k$ to the ring for a cell $k = \pi_{\text{in/out}}(j)$ are approximated by:

$$t_k = \frac{e_{\pi_{\text{in/out}}(j)} + e_{\pi_{\text{in/out}}(j+1)}}{||e_{\pi_{\text{in/out}}(j)} + e_{\pi_{\text{in/out}}(j+1)}||}, \quad n_k = \delta_{\text{in/out}}\, t_k^{\perp},$$

with $\delta_{\text{in}} = +1$ for the inner cable and $\delta_{\text{out}} = -1$ for the outer cable (if cells are ordered with increasing polar angle) and where $v^T$ denotes the transposed for any vector $v$. The orientation of the two forces are sketched in Fig. A3. Definition (11) stems from the idea that the cable acts like a wall (or an obstacle) on each particle, while definition (12) incorporates a nematic contribution [18].

#### 1.3 Scaling analysis

In this Section, we propose a dimensionless version of System (3)-(4)-(6), which we recall below:

$$\begin{aligned}
&\frac{dX_k}{dt} = V_k,\\
&V_k = cP_k + \gamma\,(F^{\text{cable}}_{\text{passive}})_k + \gamma\,(F_{\text{contact}})_k,\\
&dP_k = \mathbf{Proj}_{P_k^{\perp}} \circ \left(\nu\left(\frac{V_k}{||V_k||} - P_k\right)dt + \mu\left(\bar{P}_k - P_k\right)dt\right.\\
&\qquad\qquad \left. +\alpha_p\,(F^{\text{cable}}_{\text{passive}})_k\,dt + \alpha_a\,(F^{\text{cable}}_{\text{active}})_k\,dt + \sqrt{2D}\,(dB_t)_k\right).
\end{aligned}$$

We introduce the dimensionless variables: $\tilde{X} = X/L$, $\tilde{V} = V/c$ and $\tilde{t} = (c/L)t$, where $L$ is the characteristic length and $c$ the maximal speed. We then set this characteristic length scale to the typical length on which a cell depolarizes: $L = c/D$. We also consider re-scaled forces:

$$\tilde{F}_{\text{contact}} = \gamma/c\, F_{\text{contact}}, \quad \tilde{F}^{\text{cable}}_{\text{active}} = \beta_a\, F^{\text{cable}}_{\text{active}}, \quad \tilde{F}^{\text{cable}}_{\text{passive}} = \beta_p\, F^{\text{cable}}_{\text{passive}}.$$

where $\beta_p$ and $\beta_a$ are given force scales. All these dimensionless forces are supposed to take values of order of unity. In these new variables and dropping the "tilda" notation for simplicity, we obtain the following system:

$$\frac{dX_k}{dt} = V_k, \tag{13}$$

$$V_k = P_k + \left(\frac{\gamma\beta_p}{c}\right)(F^{\text{cable}}_{\text{passive}})_k + (F_{\text{contact}})_k, \tag{14}$$

$$dP_k = \mathbf{Proj}_{P_k^{\perp}} \circ \left(\frac{\nu}{D}\left(\frac{V_k}{||V_k||} - P_k\right)dt + \frac{\mu}{D}\left(\bar{P}_k - P_k\right)dt\right. \tag{15}$$

$$\left. +\frac{\alpha_p\beta_p}{D}\,(F^{\text{cable}}_{\text{passive}})_k\,dt + \frac{\alpha_a\beta_a}{D}\,(F^{\text{cable}}_{\text{active}})_k\,dt + \sqrt{2}\,(dB_t)_k\right). \tag{16}$$

Assuming that $\gamma\beta_p/c \ll 1$ and $\nu = \mu$, the following scaling shows that the dynamics is determined by the relative values of three parameters compared with 1:

- $\mu/D$ is the parameter of the Vicsek-like model measuring the ratio between collective alignment and noise,
- $\alpha_p\beta_p/D$ is a parameter measuring the reaction of the polarities to the cables forces,
- $\alpha_a\beta_a/D$ is a parameter measuring the active reaction of the polarities to the cables.

Note that, for the polarity dynamics, the Vicsek interactions terms (relaxation towards the averaged polarity in competition with noise) would lead to coherence as soon as the density is larger than $2D/\mu$ (in dimension 2) and to a disordered polarity distribution otherwise, as demonstrated in [6]. However, the conclusion is not valid in our case because of the boundary conditions.

### 2 Discretization, parameters and outputs

#### 2.1 Discretization

We provide in this section details about the numerical scheme implemented to solve the model given by equations (3)-(4)-(6). It is based on methods introduced in [19] for the polarity equation and [17] for the velocity equation.

We introduce a time step $\Delta t > 0$ and we aim at computing approximated positions, velocities and polarities at discrete times $t^n = n\Delta t$ with $n \in \mathbb{N}$. We denote by $(X_k^n, V_k^n, P_k^n)$ these approximated quantities. We also introduce the angle $\theta_k^n \in [0, 2\pi]$ such that $P_k^n = (\cos\theta_k^n, \sin\theta_k^n)^T$. The numerical scheme first consists into updating the polarities:

$$\begin{aligned} &\theta_k^{n+1} = \theta_k^n + 2(\hat{Q}_k^n - \theta_k^n) + \sqrt{2D\Delta t}\varepsilon_k^n, \\ &\text{with } Q_k^n = P_k^n + \frac{\Delta t}{2}\left[\left(\nu\frac{V_k^n}{\|V_k^n\|} + \mu\bar{P}_k^n + \alpha_p\,(F_{\text{passive}}^{\text{cable}})_k^n\; + \alpha_a\,(F_{\text{active}}^{\text{cable}})_k^n\right) - P_k^n\right], \end{aligned} \tag{17}$$

where for any vector $Q \in \mathbb{R}^2$, $\hat{Q}$ denotes its polar angle and where $\varepsilon_k^n$ denotes an independent realization of the standard Normal distribution. The computation of the passive force $(F_{\text{passive}}^{\text{cable}})_k^n$ is detailed in Remark 1 below. The other terms can be computed quite easily using the discrete counterparts of Equations (5) and (10)-(11)-(12).

Next, we update velocities and positions:

$$\begin{aligned} \frac{\mathbf{X}^{n+1} - \mathbf{X}^n}{\Delta t} &= \mathbf{V}^{n+1}, \\ \mathbf{V}^{n+1} &= \mathbf{Proj}_{\mathcal{C}_{\mathbf{X}^n}^{\Delta t}}\Big(c\mathbf{P}^{n+1}\Big), \end{aligned} \tag{18}$$

where the set of admissible velocities has been replaced by a first order approximation:

$$\begin{aligned} \mathcal{C}_{\mathbf{X}^n}^{\Delta t} &= \left\{\mathbf{V} \in (\mathbb{R}^2)^N \mid \quad \forall i < j, \quad D_{i,j}(\mathbf{X}^n) + \Delta t\, \nabla D_{i,j}(\mathbf{X}^n)\cdot \mathbf{V} \geqslant 0\right\} \\ &= \left\{\mathbf{V} \in (\mathbb{R}^2)^N \mid \quad \mathbf{B}\mathbf{V} - \mathbf{D} \leqslant 0\right\}, \end{aligned}$$

where $\mathbf{D} = (D_{i,j}(\mathbf{X}^n))_{i<j} \in \mathbb{R}^{\frac{N(N-1)}{2}}$, $\mathbf{B}$ is the matrix of size $(\frac{N(N-1)}{2}, 2N)$ such that $\mathbf{B}\mathbf{V} = (-\Delta t \nabla D_{i,j}(\mathbf{X}^n)\cdot\mathbf{V})_{i<j} \in \mathbb{R}^{\frac{N(N-1)}{2}}$ for any vector $\mathbf{V} \in (\mathbb{R}^2)^N$. The constraint

$\mathbf{BV} - \mathbf{D} \leqslant 0$ should be understood component-wise. In practice, this projection is reformulated as an optimization problem, that consists in finding the nearest velocity belonging to $\mathcal{C}^{\Delta t}_{\mathbf{X}^n}$:

$$\mathbf{V}^{n+1} = \underset{\mathbf{V}\in\mathcal{C}^{\Delta t}_{\mathbf{X}^n}}{\text{argmin}} \|\mathbf{V} - c\mathbf{P}^{n+1}\|^2.$$

The solution $\mathbf{V}^{n+1}$ to this minimization problem satisfies the following equations:

$$\begin{aligned}
&(\mathbf{V}^{n+1} - c\mathbf{P}^{n+1}) + \mathbf{B}^T\lambda = 0, \\
&\lambda \cdot (\mathbf{BV}^{n+1} - \mathbf{D}) = 0, \\
&\lambda \geqslant 0, \\
&\mu \cdot (\mathbf{BV}^{n+1} - \mathbf{D}) \geqslant 0, \forall \mu \geqslant 0,
\end{aligned} \tag{19}$$

where $\lambda \in \mathbb{R}^{N(N-1)/2}$ is the Lagrange multiplier to the constraints defined by the set $\mathcal{C}^{\Delta t}_{\mathbf{X}^n}$. To solve this equation, we use the Uzawa algorithm [3]. This algorithm constructs sequences $(\mathbf{V}^{(k)})_k$ and $(\lambda^{(k)})_k$ as follows:

$$\begin{aligned}
&\mathbf{V}^{(0)} = \mathbf{V}^n, \lambda^{(0)} = 0, \\
&\mathbf{V}^{(k+1)} = c\mathbf{P}^{n+1} - \mathbf{B}^T\lambda^{(k)}, \\
&\lambda^{(k+1)} = \Big(\lambda^{(k)} + \rho(\mathbf{BV}^{(k)} - \mathbf{D})\Big)_+,
\end{aligned} \tag{20}$$

where for any vector $\mathbf{u}$, $(\mathbf{u})_+ = (\max(u_i, 0))_i$ denotes the component-wise positive part of the vector and $\rho > 0$ is a numerical parameter. Under theoretical hypothesis on $\rho$, this algorithm converges: $\mathbf{V}^{(k+1)}$ and $\lambda^{(k)}$ tend to the solution $(\mathbf{V}^{n+1}, \lambda)$ of Eq. (19). For more details, we refer to [17].

To sum up, the numerical resolution of these equations is performed in four steps: first we identify the boundary cells thanks to the algorithm described in Section 1.1 and we compute the passive/active cable forces, then we compute the polarities $\mathbf{P}^{n+1}$ by (17), next the velocities $\mathbf{V}^{n+1}$ are computed thanks to the iterative algorithm (20) and finally the positions $\mathbf{X}^{n+1}$ are updated with (18).

*Remark* 1. For practical implementation, we detail here the computation of the passive force (9) given as the gradient of the elastic energy $E_{in/out}(\mathbf{X}) = E^b_{in/out}(\mathbf{X}) + E^{st}_{in/out}(\mathbf{X})$. For a given boundary cell $j' = \pi_{\text{in/out}}(j)$ of the inner or outer cable, the passive force incorporates contributions from the two springs linking it to the left and right neighboring cells, whose indexes are denoted as follows:

$$j'_p = \pi_{\text{in/out}}(j+1), \quad j'_m = \pi_{\text{in/out}}(j-1).$$

More precisely, the passive force writes:

$$(F^{\text{cable}}_{\text{passive}})_{j'} = -\nabla_{X_{j'}} E^{\text{st}}_{j'}(\mathbf{X}) - \nabla_{X_{j'}} E^{\text{st}}_{j'_p}(\mathbf{X}) - \nabla_{X_{j'}} E^{\text{b}}_{j'}(\mathbf{X}) - \nabla_{X_{j'}} E^{\text{b}}_{j'_p}(\mathbf{X}) - \nabla_{X_{j'}} E^{\text{b}}_{j'_m}(\mathbf{X}),$$

where $E^{\text{st}}_{j'}(\mathbf{X}) = k_{st}(\|X_{j'} - X_{j'_m}\| - 2r)^2$ and $E^{\text{b}}_{j'}(\mathbf{X}) = k_b(1 - e_{j'} \cdot e_{j'_p})$ are the elementary elastic energy contributions. With these notations, we have:

$$\begin{aligned}
\nabla_{\mathbf{X}_{j'}} E^{\text{st}}_{j'}(X) &= 2k_{st}(\|X_{j'} - X_{j'_m}\| - 2r)e_{j'}, \\
\nabla_{\mathbf{X}_{j'_m}} E^{\text{st}}_{j'}(X) &= -2k_{st}(\|X_{j'} - X_{j'_m}\| - 2r)e_{j'}.
\end{aligned}$$

where $e_{j'}$ was defined in (8), and

$$\nabla_{X_{j'}} E^{\mathrm{b}}_{j'}(\mathbf{X}) = k_b \begin{bmatrix} -e_{j'_p} \cdot \frac{1}{\|X_{j'}-X_{j'_m}\|} P_{e^{\perp}_{j'}} \tilde{e}_1 + e_{j'} \cdot \frac{1}{\|X_{j'_p}-X_{j'}\|} P_{e^{\perp}_{j'_p}} \tilde{e}_1 \\ -e_{j'_p} \cdot \frac{1}{\|X_{j'}-X_{j'_m}\|} P_{e^{\perp}_{j'}} \tilde{e}_2 + e_{j'} \cdot \frac{1}{\|X_{j'_p}-X_{j'}\|} P_{e^{\perp}_{j'_p}} \tilde{e}_2 \end{bmatrix}$$

$$= -\frac{k_b}{\|X_{j'} - X_{j'_m}\|} P_{e^{\perp}_{j'}} e_{j'_p} + \frac{k_b}{\|X_{j'_p} - X_{j'}\|} P_{e^{\perp}_{j'_p}} e_{j'},$$

where $\tilde{e}_1 = (1,0)^T$ and $\tilde{e}_2 = (0,1)^T$ stand for the canonical basis of $\mathbb{R}^2$. We also have:

$$\nabla_{X_{j'_m}} E^{\mathrm{b}}_{j'}(\mathbf{X}) = k_b \begin{bmatrix} e_{j'_p} \cdot \frac{1}{\|X_{j'}-X_{j'_m}\|} P_{e^{\perp}_{j'}} \tilde{e}_1 \\ e_{j'_p} \cdot \frac{1}{\|X_{j'}-X_{j'_m}\|} P_{e^{\perp}_{j'}} \tilde{e}_2 \end{bmatrix} = \frac{k_b}{\|X_{j'} - X_{j'_m}\|} P_{e^{\perp}_{j'}} e_{j'_p},$$

$$\nabla_{X_{j'_p}} E^{\mathrm{b}}_{j'}(\mathbf{X}) = k_b \begin{bmatrix} -e_{j'} \cdot \frac{1}{\|X_{j'_p}-X_{j'}\|} P_{e^{\perp}_{j'_p}} \tilde{e}_1 \\ -e_{j'} \cdot \frac{1}{\|X_{j'_p}-X_{j'}\|} P_{e^{\perp}_{j'_p}} \tilde{e}_2 \end{bmatrix} = -\frac{k_b}{\|X_{j'_p} - X_{j'}\|} P_{e^{\perp}_{j'_p}} e_{j'}.$$

#### 2.2 Parameters

The model involves 12 parameters gathered in Table A1. Whenever possible, we used physical values obtained from measurements or derived from references. It is noteworthy that only two parameters were calibrated *a posteriori*: $\alpha_p$ and $\alpha_a$ were determined by running simulations while aiming at reaching a closure time set by experiments. We provide hereafter some details on the overall procedure.

**Parameters extracted from measures.**

1. Recalling that each cell is assumed to be a rigid disk, the **diameter** $2r$ with $r$ the radius is approximated by 15 $\mu$m. This corresponds to the typical size of an undeformed cell in the initial experimental configuration.

2. We assume that the **maximal velocity amplitude** $c$ takes a value given by a normal random distribution set by experimental measurements, see Fig. S5-c. The mean value equals 21.6 $\mu\mathrm{m\,h^{-1}}$ and the standard deviation 3.7 $\mu\mathrm{m\,h^{-1}}$.

3. The **cell polarity diffusion coefficient** $D$ is extracted from individual MDCK cells, specifically from the analysis of their motion on flat surface coated with fibronectin at the same concentration as the circular patterns, see Fig. S5-de. Cell polarity orientation is set according to the cellular long axis. Its direction is defined in phase contrast by the largest lamellipodium, as shown by Fig. S5-d. Over time, cells polarities do not exhibit strong angular diffusion and three major behaviors emerge: (i) cells with low angular diffusion, (ii) cells with reversed polarity (with angle $\pi$) and (iii) cells without polarity (excluded from the analysis), see Fig. S5-d. We report the angular mean squared displacement (MSD) in Figure S5-e, yielding an angular diffusion coefficient $D = 0.016$ rad$^2$ min$^{-1}$ or 0.96 rad$^2$ h$^{-1}$.

4. To extract relevant timescales for the **relaxation parameters** $\nu$ and $\mu$, we first isolate typical examples of polarity relaxation. In mosaic experiments (see Fig. S5-f), cells with a polarity opposite to the direction of rotation assumed to be $\bar{P}_k$ have

a shift of angle $\pi$ after an average time $\tau$ and relax to $\bar{P}_k$. Consequently, considering in a first approximation that $\mu = \pi/\tau$, we obtain $\mu = 2.87 \pm 3.03$ rad h$^{-1}$ (mean $\pm$ SD). In addition, we assume that the relaxation on the velocity direction occurs with a similar timescale and thus $\nu = \mu$. In the numerical experiments, we use the value $\mu = \nu = 6.2$ rad h$^{-1}$, which gives the best fit with experiments and belongs to the considered range.

**Parameters estimated from references.**

1. The values of the **bending and stretching moduli** $k_b$ and $k_{st}$ associated to acto-myosin cables are determined from values reported for acto-myosin stress fibers.

   On the basis of a scaling argument, the stretching modulus is homogeneous to a spring constant and satisfies $k_{st} = Ed$, where $E$ is the elastic modulus expressed as a force per unit area and $d$ the thickness of the acto-myosin cable. Elastic modulus $E$ has been measured to be approximately equal to 10 kPa $= 10^4$ pN$/\mu$m$^2$ in MDCK cells [7, 24, 27] and the thickness of the acto-myosin cable $d$ approximately equals 500 nm $= 5 \times 10^{-1}\mu$m, according to [2]. Therefore, we obtain $k_{st} = 5 \times 10^3$ pN $\mu$m$^{-1}$.

   We use a similar scaling argument to derive an order of magnitude for the bending modulus $k_b$. Following [13], we consider that $k_b = \frac{\pi}{4}Ed^3$. The power $d^3$ instead of $d^4$ comes from dimensional analysis of Eq. (7). Using the previous values for $E$ and $d$, we obtain $k_b \approx 10^3$pN $\mu$m.

2. We set the **friction coefficient** $1/\gamma$ to $10^6$ pN $h\,\mu$m$^{-1}$, corresponding to a maximal friction associated with cell and/or matrix environments. We checked that the coherence and closing velocity were not significantly modified if we took cell-cell junctions friction $1/\gamma$ to $10^5$ pN $h\,\mu$m$^{-1}$ [9, 28] or cell-matrix $1/\gamma$ to $10^4$ pN $h\,\mu$m$^{-1}$ friction [15, 21].

3. Recall that the **active force** exerted on cells by the acto-myosin cables is decomposed into a symmetric and an asymmetric parts, characterized by constant amplitudes denoted by $\beta_{sym}$ and $\beta_{asym}$, respectively. In a first approximation, we assume that the two amplitudes are equal, and their value has been set to $10nN$ [25].

**Parameters obtained by calibration.**

1. **Polarizations per Newton** $\alpha_p$ and $\alpha_a$ for passive and active forces have been calibrated to recover an order of magnitude for the closing time that is similar to the experimental one. We detail this step in Section 3.1.

*Remark* 2. [Impact of parameter values on the scaling hypothesis regarding the cable forces] We recall that the passive force exerted by cables is defined in (9) and detailed in Remark 1. With the parameters of Table A1, the order of magnitude of the stretching part of the this force is $k_{st}r \approx 5 \times 10^4$ pN, while the order of magnitude of the bending part is $k_b/r \approx 10^2$ pN. Consequently, the passive force is about $5 \times 10^4$ pN and the dimensionless parameter $\gamma\beta_p/c$ is approximately $2.5 \times 10^{-3} \ll 1$. In particular, this computation justifies that the direct effect of the cable forces on the velocity dynamics can be neglected in Eq. (14).

| | | |
|---|---|---|
| Cells diameter | $2r$ | $15\ \mu$m |
| Cells maximal velocity amplitude | $c$ | $21.6\ \mu\text{m}\,\text{h}^{-1}$ (mean) |
| | | $3.7\ \mu\text{m}\,\text{h}^{-1}$ (standard deviation) |
| Angular diffusion | $D$ | $0.96\ \text{rad}^2\,\text{h}^{-1}$ |
| Relaxation parameter: polarity to velocity | $\nu$ | $6.2\ \text{rad}\,\text{h}^{-1}$ |
| Relaxation parameter: polarity to mean polarity | $\mu$ | $6.2\ \text{rad}\,\text{h}^{-1}$ |
| Cable stretching modulus | $k_{st}$ | $5 \times 10^3\ \text{pN}\,\mu\text{m}^{-1}$ |
| Cable bending modulus | $k_b$ | $10^3\ \text{pN}\,\mu\text{m}$ |
| Inverse friction coefficient | $\gamma$ | $10^{-6}\ \text{pN}^{-1}\,\text{h}^{-1}\mu\text{m}$ |
| Symmetric (wall-type) force amplitude | $\beta_{sym}$ | $10^4$ pN |
| Asymmetric (nematic-type) force amplitude | $\beta_{asym}$ | $10^4$ pN |
| Polarization per Newton parameter (passive) | $\alpha_p$ | $6.5 \times 10^{-5}\ \text{h}^{-1}\text{pN}^{-1}$ |
| Polarization per Newton parameter (active) | $\alpha_a$ | $10^{-2}\ \text{h}^{-1}\text{pN}^{-1}$ |

Table A1: (Parameters) In red: parameters extracted from measures. In blue: parameters estimated from references. In black: parameters obtained by numerical calibration.

#### 2.3 Coherence length

As introduced in the main text, the coherence length measures the level of synchronization of cells in rings. We denote by $v(\theta)$ the tangential velocity in polar coordinates. The coherence length is then defined by:

$$\xi = -\frac{1}{2\pi}\frac{C_v(\theta)}{C_v(0)}$$

where $C_v$ denotes the correlation of the tangential velocity:

$$\mathrm{C}_v(\theta) = \int_0^{2\pi} v(t+\theta)v(t)dt.$$

In practice the correlation is computed using its expression:

$$\mathrm{C}_v(\theta) = \mathcal{F}^{-1}(|\mathcal{F}(v)|^2)(\theta).$$

where $\mathcal{F}$ and $\mathcal{F}^{-1}$ denotes the Fourier transform and its inverse (well defined on smooth functions). We note that this quantity is rescaled by the length of the circle: $\xi$ near 1 indicates that the characteristic length scale is equal to that of the circle and thus corresponds to full coherence. As a larger characteristic length scale also indicates full coherence, we consider a truncated version of this quantity: $\min(\xi, 1)$.

We use this indicator on the tangential polarity density $v(\theta)$. We compute this quantity at the discrete level by using a frame centered at the barycenter of the cell positions and sampling the domain in the polar direction with a fixed number of circular sectors $\theta_j = 2\pi j/N_\theta$ for $j \in \{0, N_\theta - 1\}$ with $N_\theta = 24$ similar to experiments and we write $d\theta = 2\pi/N_\theta$. For each cell, we denote by $(r_k, \theta_k)$ its polar coordinates in this frame. Then we compute:

$$v(\theta_j) = \frac{\frac{1}{N}\sum_{k=1}^N K(\theta_k - \theta_j)V_k \cdot e_{\theta_k}}{\frac{1}{N}\sum_{k=1}^N K(\theta_k - \theta_j)}, \qquad \forall j \in \{0, N_\theta - 1\},$$

where $V_k \cdot e_{\theta_k}$ denotes the tangential velocity of the $k$-th cell, the kernel function $K(\theta) = \mathbf{1}_{|\theta|<d\theta}|\theta/d\theta|/d\theta$ represents the density contribution of a cell at distance $\theta$ from 0 and $\mathbf{1}_{|\theta|<d\theta}$ is the characteristic function of the set $[-d\theta, d\theta]$, taking the value 1 inside the domain and vanishing elsewhere. We thus obtain a discrete representation $(v(\theta_j))$ of the tangential velocity density $v(\theta)$. Finally we compute the coherence length by first computing the correlation with the discrete Fourier transform and then using a discrete finite difference formula.

### 3 Simulation set up and numerical experiments

We consider a ring domain $\Omega = \{(r, \theta) \in [R_{\min}, R_{\max}] \times [0, 2\pi[\}$ and run simulations in the following four experimentally guided configurations: $R_{\min} = 40, 90, 150$ or $500$ $\mu$m. In each case, $R_{\max} = R_{\min} + 30$ $\mu$m, corresponding to the rings of (internal) diameter 80 $\mu$m, 180 $\mu$m, 300 $\mu$m and 1000 $\mu$m respectively. As cells have diameter equal to 15 $\mu$m, two cells can be present on each radial section of the annular at initial condition. Accordingly, by taking into account the annular radius, the number of cells is set to 46, 89, 140 or 439, respectively.

Initially, cells are distributed uniformly on the annular domain $\Omega$. The influence of the initial polarity distribution will be assessed by means of numerical experiments in Sec. 3.3. We assume that velocities are initially aligned with the polarities up to the volume-exclusion constraint. The values used for the parameters are provided in Sec. 3.1 and the numerical experiments performed in the calibration procedure are detailed in the next subsections.

In all simulations, the time step is chosen equal to $\Delta t = 10^{-3}$h and the final time $T = 24$h. Time step is chosen such that the dynamics is well resolved in time. As regards to the Uzawa algorithm for computing the volume-exclusion force, we use a stopping criteria on the relative distance between two iterative values of the velocity field with a threshold value of $10^{-3}$ and $\rho$ is taken equal to $(12\sqrt{2}\Delta t^2)^{-1}$ to ensure convergence.

#### 3.1 Calibration of the polarization coefficient

To calibrate the parameter $\alpha_p$ involved in Equation (6), we focus on the experimental closing time of the 80 $\mu$m rings. In Figure A4 (left) obtained with $\alpha_a = 10^{-2}$ h$^{-1}$pN$^{-1}$, we display results of simulations corresponding to values $\alpha_p$ in $\{5 \times 10^{-5}$ h$^{-1}$pN$^{-1}, 6 \times 10^{-5}$ h$^{-1}$pN$^{-1}, 6.5 \times 10^{-5}$ h$^{-1}$pN$^{-1}, 7 \times 10^{-5}$ h$^{-1}$pN$^{-1}, 8 \times 10^{-5}$ h$^{-1}$pN$^{-1}, 1 \times 10^{-4}$ h$^{-1}$pN$^{-1}\}$. We assess the impact of this parameter which is the main one driving the closing time scale and we select the value $6.5 \times 10^{-5}$ h$^{-1}$pN$^{-1}$ in order to retrieve the experimentally reported closing time. In Figure A4 (right), we also observe that this choice has no significant influence on the coherence acquisition for both velocity fields.

We emphasize that, when considering rings of diameter 180 $\mu$m, the same values for this parameter give consistent results with the experimental dynamics (see Figure A5). Thus this numerical experiment can be seen as a further validation for this calibration procedure.

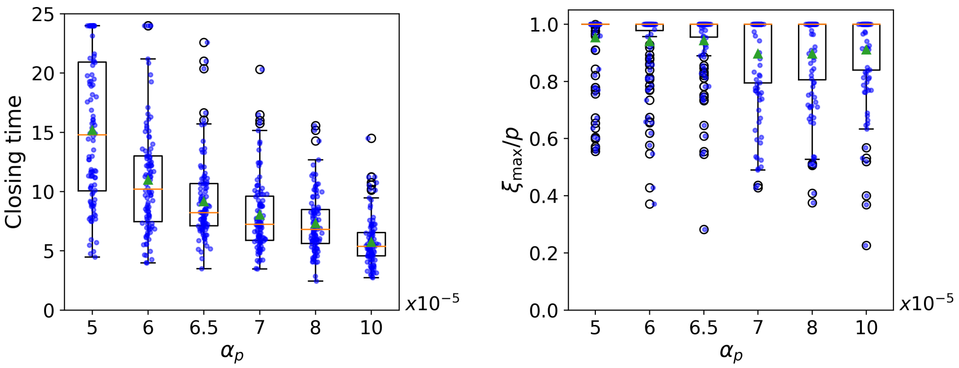


Figure A4: Effect of the variation of the parameter $\alpha_p$ on (i) the closing time; (ii) the maximal coherence length of the velocity field. The median value is represented as an orange horizontal line, and the mean depicted as a green triangle.

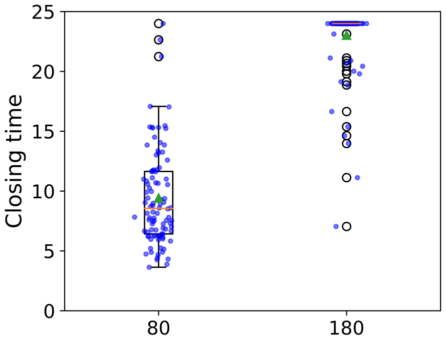


Figure A5: Closing time for rings of diameter 80 and 180 $\mu$m. The median value is represented as an orange horizontal line, and the mean depicted as a green triangle.

#### 3.2 Influence of the distribution of cells maximal velocity amplitude

Contrary to the other parameters which are fixed, the cells maximal velocity amplitude $c$ is selected for each simulation following a normal distribution of mean 21.6 $\mu$m h$^{-1}$ and standard deviation 3.7 $\mu$m h$^{-1}$. To assess the influence of this assumption on our findings, we compare in Figure A6 results obtained with normal distribution and with a constant $c = 21.6\,\mu$m h$^{-1}$. The qualitative variation of the coherence length with respect to the diameter of the rings with normal distribution provides good agreement with the experimental data compared to the constant case, which validates our assumption.

#### 3.3 Effects of the coherence length

Results displayed in Fig.5-d show that the average time to acquire coherence is larger when the initial coherence length is small. As a complement, we display in Figure A7 the closing time (left panel) and the maximal coherence length during the entire simulation (right panel) as functions of the initial coherence length. As expected, the larger the initial coherence, the larger the closing time, as the coherence counterbalances the radial

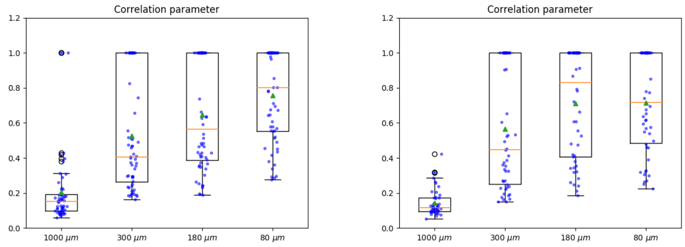


Figure A6: Comparison of coherence length with the choice of cells maximal velocity amplitude. Left: $c$ selected following a normal distribution of mean $21.6\,\mu\text{m}\,\text{h}^{-1}$ and standard deviation $3.7\,\mu\text{m}\,\text{h}^{-1}$. Right: constant $c = 21.6\,\mu\text{mh}^{-1}$. The median value is represented as an orange horizontal line, and the mean depicted as a green triangle.

bending forces. Likewise, an initial coherence larger than 0.4 leads systematically to coherence acquisition, whereas smaller ones induce more scattered behaviour.

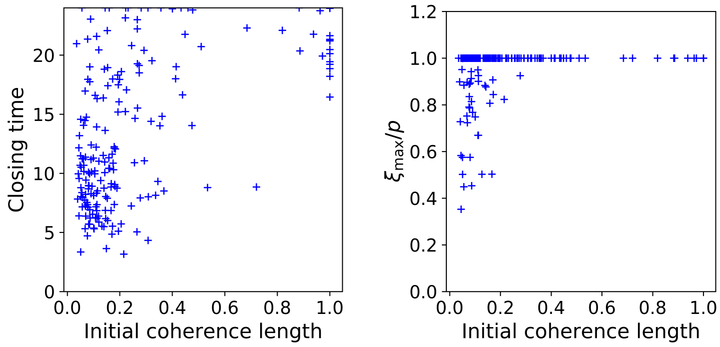


Figure A7: Variation of closing time (left) and the maximal coherence length on the whole simulation (right) with respect to the initial coherence length.

These results suggests that there may be a critical value of coherence length that induces the onset of coherence. In Figure A8 (left), we plot the probability to reach coherence after having reached a given coherence length. This has been obtained using 200 different simulations with rings of diameter $80\,\mu$m and parameters of Table A1. We observe that above a coherence length equal to 0.75, the probability to reach coherence significantly increases. In Figure A8 (right), we depicted the time evolution of 50 different numerical simulations to illustrate the stochastic variability of the model. The dynamics are colored in blue after the closing time. On each plot, marks have been added to identify the first time a coherence length equal to 0.75 is reached (red point), and the first time the system reaches full coherence (blue point).

Coherence length has been so far chosen as the indicator to characterize the onset of coherence. Another possible indicator would be the percentage of cells with a clockwise-oriented velocity in the ring. Indeed, coherence is also associated to a large percentage of aligned cells. However, these two indicators are not identical: Fig. A9 (left) actually shows that a similar percentage of oriented cells can be associated with very different values of coherence length. On Fig. A9 (right), we look at the probability to reach coherence after having reached a percentage of aligned cells. It seems that the critical value of 0.75 can also be an indicator for the onset of coherence.

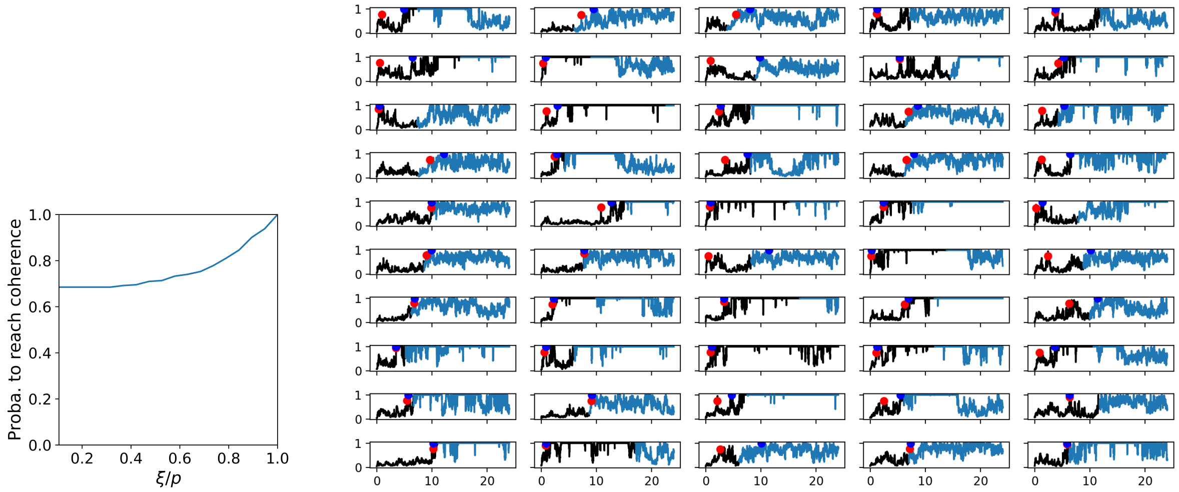


Figure A8: Left: Probability (percentage of numerical simulations) to reach coherence after having reached a given coherence length $\xi/p$. Right: Coherence length over time for 50 numerical simulations. For each simulation, the dynamics are colored in blue after the closing time. The red point (resp. blue point) correspond to the first occurrence of the coherence length equal to 0.75 (resp. 1).

#### 3.4 Effect of the inner/outer cables

Numerical results displayed in Fig.5 demonstrate the importance of the contribution of active forces at cables, which have a passive component corresponding to the stretching and bending moduli of the acto-myosin structure, and an active component translating the contribution of cables in setting tangential polarities. The active component was required for the onset of coherence (Fig. 5e). We further investigated the effect of the properties of the cables in two experimentally meaningful configurations: (i) the Caldesmon experiment (Fig. 3f) and (ii) the experiment on cell migration confined in a ring-shaped pattern described in [12].

In the first configuration (i), we tested the effect of introducing a cell with no cable within the ring, thereby modelling the CaD mosaic rings experimental setting (Fig. 3f). Results displayed in Fig. 5f and Movie S14 show that the phenotype of this cell escape is reproduced, substantiating the confinement role of cables.

A second numerical experiment (ii) was designed to simulate a confined environment [12], by changing the properties of the actomyosin cables to make them very stiff ($k_{st}^{\text{stiff}} = 10^5 \times k_{st}$, $k_b^{\text{stiff}} = 10^5 \times k_b$ and $\alpha_p$ in Eq. (16) scaled accordingly). Note that the scaling analysis from Sec. 1.3 (see also Remark 2) shows that in this case, the order of magnitude of $\gamma\beta_p/c$ is not negligible anymore and therefore the contribution of the term pertaining to the cable forces on the velocity dynamics is accounted for in Eq. (14). We also deactivate the dynamical update of the cells involved in the cables to prevent the ring from contracting when the number of cells per cables decreases, and thus to preserve the geometry of the ring over time. In Fig. A10, we performed a similar numerical experience as in Fig. A8: in contrast to the case of elastic cables considered in the present work, the coherent motion occurs systematically for confined rings of diameter 80 $\mu$m. Note however that these simulations are not strictly equivalent to a confined environment dynamics as

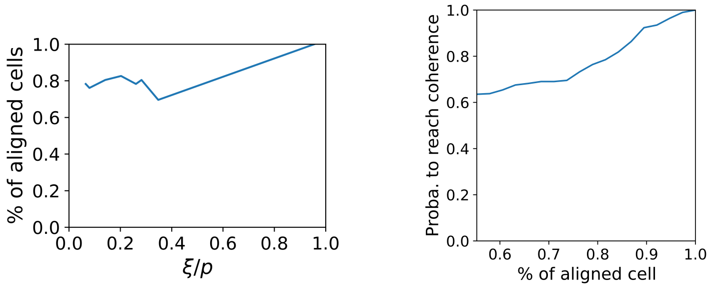


Figure A9: Left: Example of percentage of aligned cells as a function of the coherence length for different initial velocity configurations. The initial tangential velocity is here taken piece-wise constant, equal to one on periodic intervals (aligned zone) and zero elsewhere (non-aligned zone). Right: Probability (percentage of numerical simulations) that reach coherence after having reached a given percentage of aligned cells.

collective translation of the ring is possible.

We also investigated the dynamics of larger rings of diameter 180 $\mu$m, 300 $\mu$m and 1000 $\mu$m respectively. As their smaller curvatures induce smaller bending forces, we have to consider stiffer cables to make the passive cable force effective for the two larger diameters. Remarkably, as shown in Figure A11, the minimal model we designed is able to systematically reproduce the onset of a coherent motion in all situations without any other necessary changes.

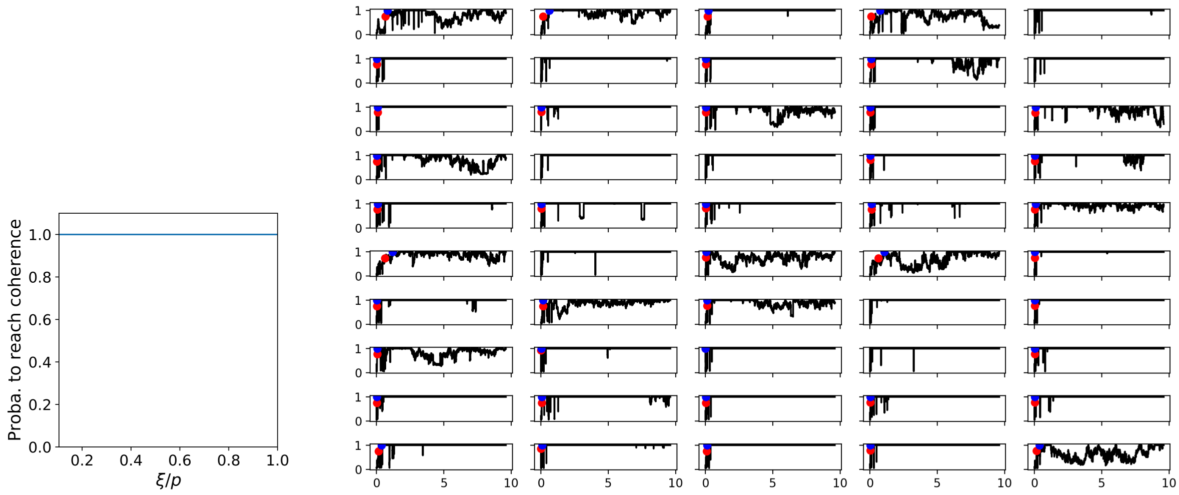


Figure A10: Dynamics with stiff cables. Left: Probability (percentage of numerical simulations) to reach coherence after having reached a given coherence length $\xi/p$. Right: Coherence length over time for 50 numerical simulations. The red point (resp. blue point) correspond to the first occurrence of the coherence length equal to 0.75 (resp. 1).

#### 3.5 Phase diagrams

The aim of this section is to numerically explore the complex behavior of the 80 $\mu$m diameter rings and the onset of a coherent motion *via* a thorough analysis of dynamic

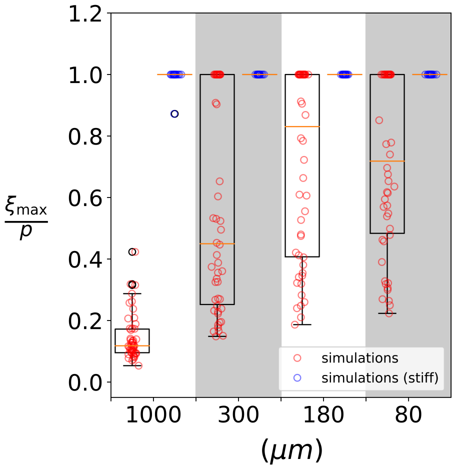


Figure A11: Comparison of maximal coherence length obtained for simulations with physical cables (in red) and with stiff cables (in blue) for different ring diameters.

phase diagrams. A first approach would have been to study coherence as a function of cell polarity and velocity/RhoA activity. Nevertheless, the theoretical model involves 12 parameters (see Sec. 3.1) and it would have been difficult to single out each contribution with this direct approach. We therefore performed in Sec. 1.3 a scaling analysis and thus focused our exploration of the parameter space on the newly identified 3 meaningful values of $\mu/D$, $\alpha_a\beta_a/D$, $\alpha_p\beta_p/D$, respectively.

In Fig. A12 below, we first plot the phase diagram when $\alpha_p\beta_p/D$ is kept fixed. We thus measure the behavior of the system according to the two active components of the dynamics: the collective alignment term and the active reaction to the cables. On each plot, the solid middle surface corresponds to the average values of 50 simulations for each set of parameters. In the top left panel, we observe that for active systems with large $\alpha_a\beta_a/D$, the closure time of the ring is larger, suggesting that active term related to the cable slows down the closing of the ring. Regarding the maximal coherence length over the simulations, the top right panel shows that it increases with the level of collective alignment dynamics. In the bottom left panel, the tangential velocity order parameter computed is displayed. It increases with both activity terms. This means that an increased of order parameter is associated with the onset of the collective behaviour. Finally, the bottom right panel shows the dimensionless maximal diameter (maximal diameter divided by the initial one, which equals 140): a larger cable activity leads to an increase of the maximal radius of the rings and may be far from the experimental observations.

Next, Fig. A13 presents the phase diagram when $\mu/D$ is fixed and we study both the influence of the active and passive reaction of the polarities to the cable. First, the top left panel shows that the active reaction makes the closure time increase while the passive term makes it decrease: when polarities aligns to the cable force, this prevents the ring to extend too much. Indeed, the bottom right panel indicates that except for large active and small passive parameters, the ring is actually extending and the model is less relevant as this implies long range interactions between cells. On the top right and bottom left panels, we observe that the maximal coherence length and the tangential velocity order

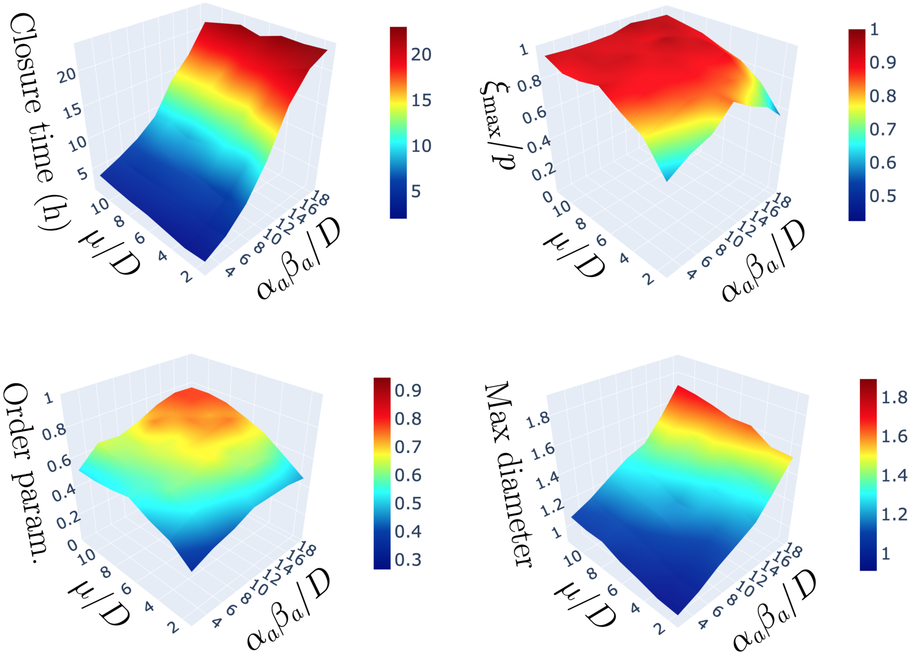


Figure A12: Closure time (time in hours (h), top left panel), maximal coherence length (dimensionless, top right panel), order parameter (dimensionless, bottom left panel) and maximum diameter (re-scaled by the initial diameter of the ring, top right panel) as functions of the dimensionless parameters $\mu/D$ and $\alpha_a\beta_a/D$. On each plot, each point of the surface corresponds to the average value of 50 simulations.

parameter have weak variations on the consider range of parameters. Combined with the phase portraits on Fig. A12, we may conclude that coherence is strongly dependent on the fixed collective alignment parameter $\mu/D$.

#### 3.6 Performance of the minimal model and limitations

The theoretical formulation of the model is quite generic. In particular, we focused in the present contribution on two-dimensional case since the experimental dynamics is also mainly two-dimensional. However cells width could have been considered with associated 3D simulations. This was not necessary in the present contribution since we show that key features were already be captured in two dimensions.

Cell deformation and cell division were not incorporated in the model. However with our simple rigid particles, simulations were able to successfully reproduce the main features of experimental dynamics. Further extension of the model could include the cell deformability (along [22]) and cell division [29].

Regarding the description of cable and its impact on the overall dynamics, we do not account for the formation of holes, since more elaborate algorithms would be required for topological changes. In what concerns the description of the forces acting on the cells at the edge of the cell layer, the idea of incorporating the restoring force due to curvature, the active lamellipodia-driven force acting on the convex regions and the force arising from the contractile acto-myosin cable acting in the concave regions could have been considered, as in [29]. As an alternative, a macroscopic description by means of level-set

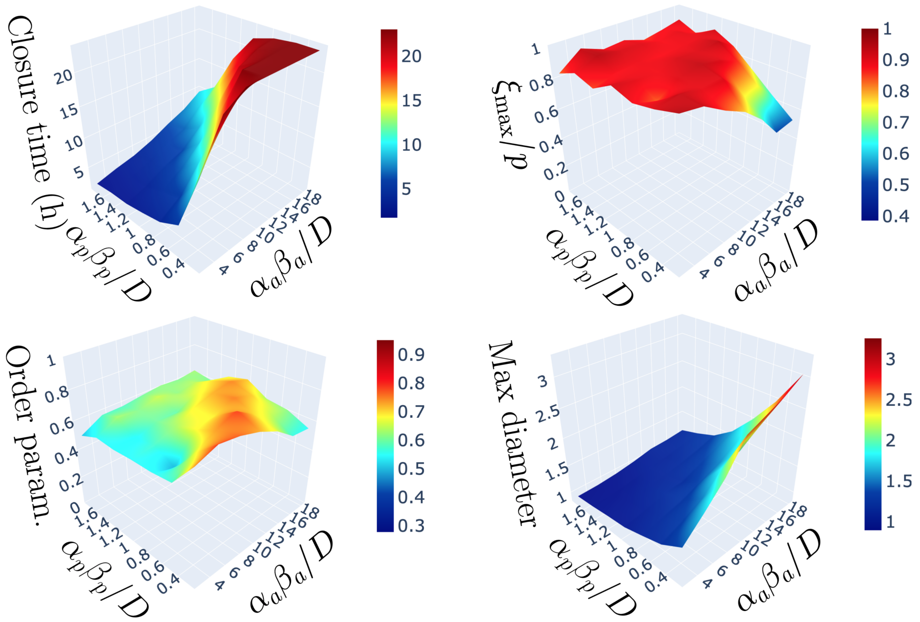


Figure A13: Closure time (time in hours (h), top left panel), maximal coherence length (dimensionless, top right panel), order parameter (dimensionless, bottom left panel) and maximum diameter (re-scaled by the initial diameter of the ring, top right panel) as functions of the dimensionless parameters $\alpha_a\beta_a/D$ and $\alpha_p\beta_p/D$. On each plot, each point of the surface corresponds to the average value of 50 simulations.

functions could be envisioned [8].

More generally, from a theoretical perspective and in order to answer to the challenging question of the possible connection between the agent-based model on the one hand and a continuum-level description on the other hand, it would be interesting to derive a macroscopic limit of the model in the spirit of [4]. At the macroscopic level, an alternative approach would be to use active nematic hydrodynamics models [1, 30]. However, establishing the link between particle descriptions and such models is still challenging from a mathematical point of view.
